# Substrate Transport and Specificity in a Phospholipid Flippase

**DOI:** 10.1101/2020.06.24.169771

**Authors:** Yong Wang, Joseph A Lyons, Milena Timcenko, Felix Kümmerer, Bert L. de Groot, Poul Nissen, Vytautas Gapsys, Kresten Lindorff-Larsen

## Abstract

Type 4 P-type ATPases are lipid flippases which help maintain asymmetric phospholipid distribution in eukaryotic membranes by driving unidirectional translocation of phospholipid substrates. Recent cryo-EM and crystal structures have provided a detailed view of flippases, and we here use molecular dynamics simulations to study the first steps of phospholipid transport and lipid substrate specificity. Our simulations and new cryo-EM structure shows phospholipid binding to a groove and subsequent movement towards the centre of the membrane, and reveal a preference for phosphatidylserine lipids. We find that only the lipid head group stays in the groove while the lipid tails remain in the membrane, thus visualizing how flippases have evolved to transport large substrates. The flippase also induces deformation and thinning of the outer leaflet facilitating lipid recruitment. Our simulations provide insight into substrate binding to flippases and suggest that multiple sites and steps in the functional cycle contribute to substrate selectivity.

## Introduction

Eukaryotic cells separate the inside of the cell from the external environment by a complex mixture of phospholipids (PLs), proteins and other molecules which together form a ∼4 nm thick bilayer (***Kobayashi and Menon, 2018***). The outer and inner leaflets of plasma membranes have different lipid composition, saturation and packing (***Kobayashi and Menon, 2018; Levental et al., 2020***), so that e.g. phosphatidylserine (PS) and phosphatidylethanolamine (PE) are enriched in the inner (cytosolic) leaflet, while phosphatidylcholine (PC) and sphingomyelin are found more often in the outer (exoplasmic) leaflet. The maintenance or modulation of the asymmetric PL distribution can play a central role in the proper functioning of the cell; for example, an increase in the population of PS in the outer leaflet of the plasma membrane serves as an ‘eat-me’ signal to induce phagocytosis (***Fadok et al., 1992***), whereas a decrease of the population of the negatively charged PS on the inner leaflet may result in a release of proteins with positively charged residues from the membrane to the cytosol by an ‘electrostatic switch’ mechanism (***Lipp et al., 2019***).

It is kinetically unfavourable for PLs to ‘flip’ (move from the outside to inside) and ‘flop’ (inside to outside), and it requires energy to maintain asymmetric lipid distributions. Therefore, cells contain a number of different PL transporters that fall into three broad categories: scramblases, floppases, and flippases (***Montigny et al., 2016***). Scramblases (such as TMEM16 (***Falzone et al., 2018***) and a number of G-protein coupled receptors (***Menon et al., 2011***)) facilitate the movement of PLs in both directions without energy consumption. In contrast, flippases and floppases are unidirectional and ATP-dependent. They use the energy from ATP hydrolysis to flip or flop specific PLs against their concentration gradient. Indeed, PL flip/flop in the presence of ATP-dependent flippases/floppases typically occurs at the time scale of ∼10-100 ms, much faster than that of the spontaneous process which is typically several hours or longer (***Kobayashi and Menon, 2018***).

The type 4 (P4) subgroup of the P-type ATPase superfamily are lipid flippases, that are related to other well-characterized P-type ATPases including the Na^+^/K^+^-ATPase (NKA) and sarco-endoplasmic reticulum Ca^2+^-ATPase (SERCA) of the type II subgroup (***Palmgren and Nissen, 2011; Andersen et al., 2016***). NKA maintains the electrochemical gradients for Na^+^ and K^+^ (***Toyoshima et al., 2011***), while SERCA actively pumps Ca^2+^ from the cytosol into internal stores (***Carafoli, 2002***). P-type ATPases are predicted to share a common core topology of cytosolic A (actuator), P (phosphorylation), and N (nucleotide binding) domains, which confer the ATPase functionality, and a transmembrane (TM) domain (***Palmgren and Nissen, 2011***).

Transport by P-type ATPases generally follows the so-called Post-Albers cycle (***Albers, 1967; Post et al., 1972***), which has been structurally described by a wealth of crystal structures of SERCA and NKA (***Dyla et al., 2020***). The cycle transitions between two main functional states (E1 and E2) and their phosphorylated forms (termed E1P and E2P) (Fig. 1). During each cycle, inward- and/or outward-transport is controlled by opening and closing of cytoplasmic and exoplasmic pathways, which give access to the transmembrane binding sites with distinct affinities and specificities for the transported substrates. The transport of lipids by P4-ATPases is associated with the dephosphorylation reaction, i.e., the E2P to E2/E1 transition (***Lyons et al., 2020***). Given the fact that PLs are typically ∼10-fold larger than the cations transported by the P2 ATPases, the question, also referred to as the ‘giant substrate problem’ (***Baldridge and Graham, 2012***), arises as to how P4 ATPases could use the same ‘alternating-access’ mechanism to transport large PLs.

**Figure 1.**
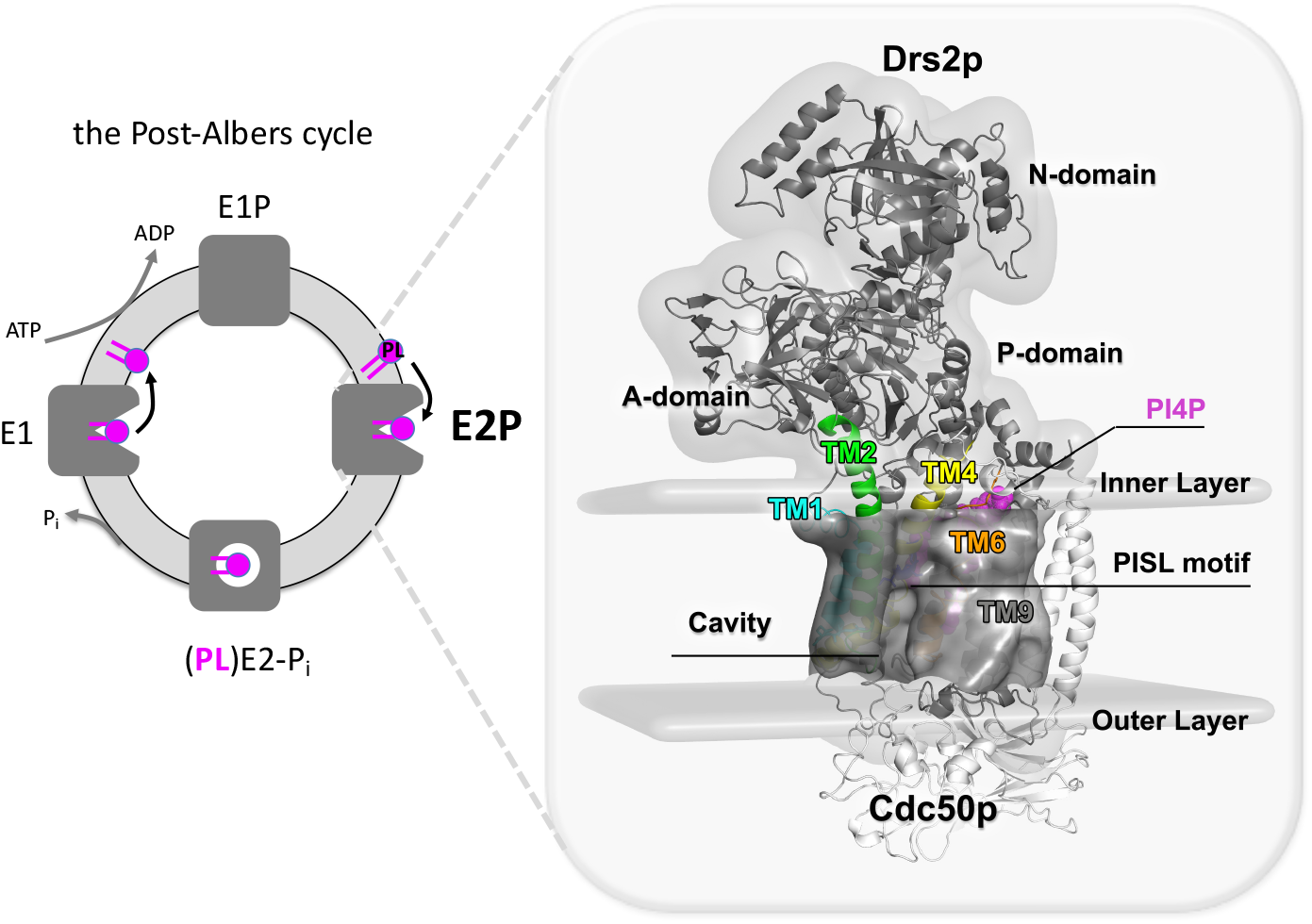
The general (Post-Albers) conformational cycle of flippases and the cryo-EM structure of PI4P-activated E2P state of Drs2p in complex with Cdc50p. The left panel shows a simplified Post-Albers transport cycle for P4-ATPases, in which repeating conformational changes occur between a few key functional states: the ‘inward-open’ E1, the phosphorylated E1P, the ‘outward-open’/occluded E2 and E2P. The yeast lipid flippase Drs2p has an architecture similar to that of other P-type ATPases, consisting of an actuator (A) domain, a nucleotide-binding (N) domain, a phosphorylation (P) domain, and a transmembrane domain of ten transmembrane helixes (TMs). Drs2p functions in the presence of an auxiliary subunit, Cdc50p, which may play a role as a chaperone via an unknown mechanism. The Drs2p–Cdc50p complex alone is in an inactive form autoinhibited by the C-terminal tail of Drs2p (not shown in the cycle), but is activated by the binding of lipid phosphatidylinositol-4-phosphate (PI4P). The right panel shows the cryo-EM structure of the Drs2p in complex with Cdc50p (white cartoon) in the PI4P-activated E2P state (PDB entry: 6ROJ) (***Timcenko et al., 2019***).

The first structural insights into P4 flippase function were recently provided by the cryogenic electron microscopy (cryo-EM) structure of a key functional E2P state of the yeast Drs2p P4-ATPase (***Timcenko et al., 2019***). Similar structures of Drs2p and structures of other flippases, including mammalian ATP8A1, ATP11C and yeast Neo1p, Dnf1p and Dnf2p have been also subsequently determined (***Bai et al., 2019; Hiraizumi et al., 2019; Nakanishi et al., 2020a,b; He et al., 2020; Bai et al., 2020, 2021; Timcenko et al., 2021***). Drs2p is found in the trans-Golgi network membrane, where it acts as a PS lipid flippase, but has also been shown to transport PE to a lesser extent (***Natarajan et al., 2004; Azouaoui et al., 2014***). Like Drs2p, ATP8A1 and ATP11C also preferentially transport PS and PE, whereas Dnf1p and Dnf2p prefer PE, PC and their lyso-forms, as well as monosaccharide glycosphingolipids (like glucosylceramide and galactosylceramide) (***Baldridge and Graham, 2012; Takatsu et al., 2014; Lee et al., 2015; Roland et al., 2019***). The published structures provide molecular details on the transport cycle and lipid translocation, revealing putative binding locations on the protein surface for substrate entry (Fig. 1). These stabilized structures, however, do not directly reveal the process by which lipids are recognized and transported, and detailed analysis is further hampered by the lack of clear density and unambiguous assignment for the substrates in most of the structures.

Prior to the determination of atomic structures, a few models had been proposed to describe the mechanism of substrate selectivity. The ‘credit card’ model, inspired by scramblases (***Pomorski and Menon, 2006; Morra et al., 2018***), proposes a peripheral pathway that allows for lipid translocation. Here, the lipid flippase recognises and occludes the lipid head group while the hydrophobic tails remain in the membrane. Consistent with the credit card hypotheses, three transport models identifying different lipid entry sites on the protein surface have been proposed, namely the two gate, hydrophobic gate and central cavity models. The ‘two-gate’ model proposed the presence of an entry gate where the PL is initially selected from the outer leaflet by the residues clustering at an entry site defined between TM1, TM3 and TM4 and an exit gate on the inner leaflet side of the membrane for further substrate selection (***Baldridge and Graham, 2013***). The ‘hydrophobic gate’ model derived from mutagenesis of bATP8A2 and a homology model based on SERCA, suggested that a ring of hydrophobic residues centred around the highly conserved residues in the unwound section of TM4 (PISL in the case of Drs2p) separates two proposed water-filled cavities that facilitate lipid transport via a pathway defined by TM1, TM2, TM4 and TM6 (***Vestergaard et al., 2014***). The ‘central cavity’ model based on mutagenesis data and homology modeling proposed that a large and deep cavity formed in the center of the TM domain bordered by TM3, TM4, and TM5 could potentially accommodate the PL headgroup during transporting across the membranes analogous to the ion transport mechanism in SERCA and NKA (***Jensen et al., 2017***). These models alone, however, do not explain all of the biochemical data (***Andersen et al., 2016; Roland and Graham, 2016; Palmgren et al., 2019***). The recent substrate-bound flippase structures are more consistent with the ‘hydrophobic gate’ model. The structural determinants of lipid specificity and recognition, however, remain elusive.

Here, we use MD simulations in both atomic and near-atomic detail to explore the initial lipid binding process in the outward-open E2P state and investigate the substrate selectivity using alchemical free energy calculations. We also present the cryo-EM structure of a substrate-bound E2P state of Drs2p and perform atomistic MD simulations of this state, that together reveals specific interactions between the lipid head group and the flippase. Our results reveal a preference for PS binding to a water-filled substrate-binding groove, even in the outward open conformation, and local deformations and thinning of the lipid bilayer around Drs2p.

## Results

### Phospholipid binding to the TM2-TM4-TM6 groove of Drs2p

We used a combination of all-atom and coarse-grained (CG) MD simulations to investigate the lipid binding preferences of Drs2p, and how these depend on the lipid composition and membrane environment. Thus, we inserted the EM structure of the PI4P-activated E2P state of the Drs2p-Cdc50p complex into a series of simplified lipid bilayer models consisting of POPS (1-palmitoyl-2-oleoylsn-glycero-3-phosphoserine), POPE (1-palmitoyl-2-oleoyl-sn-glycero-3-phosphoethanolamine) and POPC (1-palmitoyl-2-oleoyl-sn-glycero-3-phosphocholine). The bilayers are asymmetric and have POPE and POPS in the inner leaflet at a 2:1 ratio, and POPS, POPE and POPC in the outer leaflet at different ratios. To mimic a biologically more realistic membrane similar to that found in yeast (***Daum et al., 1999***), we also inserted the system in a bilayer consisting of seven different lipid types: POPS, POPE, POPC, POPA (phosphatidic acid), POPI (phosphatidylinositol), POPG (phosphatidylglycerol) and ERG (ergosterol) at a 2:10:15:2:10:1:10 ratio. All together we simulated the behaviour of Drs2p in six atomistic and eight CG models (Table 1; Materials and Methods).

**Table 1.** Atomistic and CG MD simulations of PI4P activated E2P state of the Drs2p-Cdc50p complex.

| Name | Model | Components of luminal leaflet | Length |
| --- | --- | --- | --- |
| CG1 | Martini2.2 | 100%POPS | 2x100 μs |
| CG2 | Martini2.2 | 100%POPE | 2x100 μs |
| CG3 | Martini2.2 | 100%POPC | 2x100 μs |
| CG4 | Martini2.2 | 50%POPE:50%POPS | 2x100 μs |
| CG5 | Martini2.2 | 33.3%POPC:33.3%POPE:33.3%POPS | 3x100 μs |
| CG6 | Martini2.2 | 50%POPC:50%POPS | 2x100 μs |
| CG7 | Martini2.2 | 50%POPC:50%POPE | 2x100 μs |
| CG8 | Martini2.2 | 1POPS:1POPE:1POPC:1POPG:1POPA:1POPI:2CHOL | 4x10 μs |
| AA1 | CHARMM36m | 100%POPS | 500 ns |
| AA2 | CHARMM36m | 100%POPE | 500 ns |
| AA3 | CHARMM36m | 100%POPC | 500 ns |
| AA4 | CHARMM36m | 50%POPE:50%POPS | 500 ns |
| AA5 | CHARMM36m | 33.3%POPC:33.3%POPE:33.3%POPS | 3+2+2 μs |
| AA6 | CHARMM36m | 4%POPS:20%POPE:30%POPC:2%POPG:4%POPA:20%POPI:20%ERG | 500 ns |

In the atomistic simulations, we observed rapid (within tens of nanoseconds) binding of lipids to the entry of a groove formed between TM2, TM4 and TM6 (Fig. 2A and B). We observed that the phosphate headgroup of the bound lipids moved ∼0.7-1.0 nm inside the membrane (Fig. 2A), and found that all three lipids (POPS, POPE and POPC) could bind to a similar location in different outer leaflet models. In our simulations of the yeast-like membrane model, we did not observe any substantial differences in the behaviour of protein and other related properties (such as the membrane deformation, water cavities, etc.) as compared with the simulations in the PS/PE/PC models. Therefore, to facilitate the investigations on the lipid competition and specificity, we focused on the results of the simplified models composed of PS/PE/PC unless otherwise specified.

**Figure 2.**
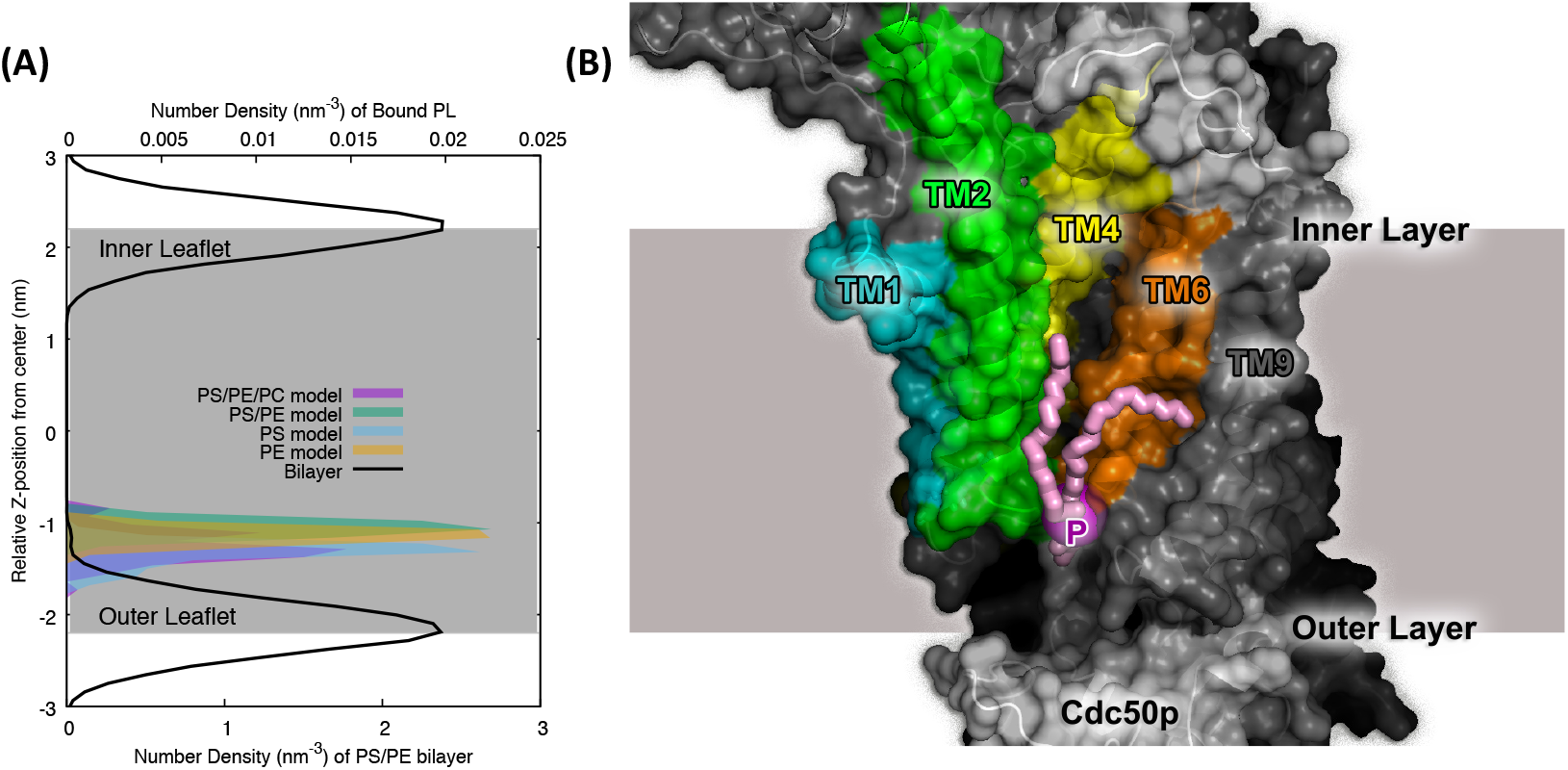
Initial phospholipid binding to Drs2p observed in atomistic simulations. (A) The average density maps of the phosphate headgroup of the bound PLs along the Z axis relative to the membrane center (z=0) obtained from the last 100 ns of the atomistic simulations with different outer leaflet models. The density profile of the lipid bilayer (black line) is shown for comparison. (B) The representative lipid bound conformation, shown using a snapshot from an atomistic simulation of the pure POPS outer leaflet model (AA1; see also Table 1). TM1, TM2, TM4, TM6 are colored in cyan, green, yellow, and orange, respectively. The phosphorus atom of the bound PL is represented by a magenta sphere and the acyl chains are represented by pink sticks. The membrane bilayer is shown as a grey background.

As our atomistic simulations were limited to a few *µs*, they did not allow us to observe lipid exchange and competition within the groove, nor to investigate whether the substrate can bind more deeply inside the protein. We therefore resorted to CG modeling using the popular Martini model (***Marrink et al., 2007***) which has been successfully applied to study a wide range of membrane protein systems and their protein-lipid interactions (***Corradi et al., 2018, 2019; Muller et al., 2019; Ansell et al., 2021***). For each lipid and bilayer model, we performed two (replica) simulations, each 100 *µs* long, (we included an extra replica for the POPS/POPE/POPC mixed outer leaflet model), resulting in 1.5 ms of simulations in total, which enabled us to observe additional lipid binding events and better sample protein-lipid interactions. We calculated the average density map of the phosphate headgroup of the lipids which revealed clusters of lipid binding sites (Fig. 3A). In addition to multiple lipid binding sites on the protein surface, we found substantial lipid binding in the TM2-TM4-TM6 groove. This occurs both at the mouth of the groove, similar to what we observed in the atomistic simulations, and deeper inside the groove all the way up to the conserved PISL motif that has been speculated to act as a gate between entry and exit sites (***Vestergaard et al., 2014; Timcenko et al., 2019***).

**Figure 3.**
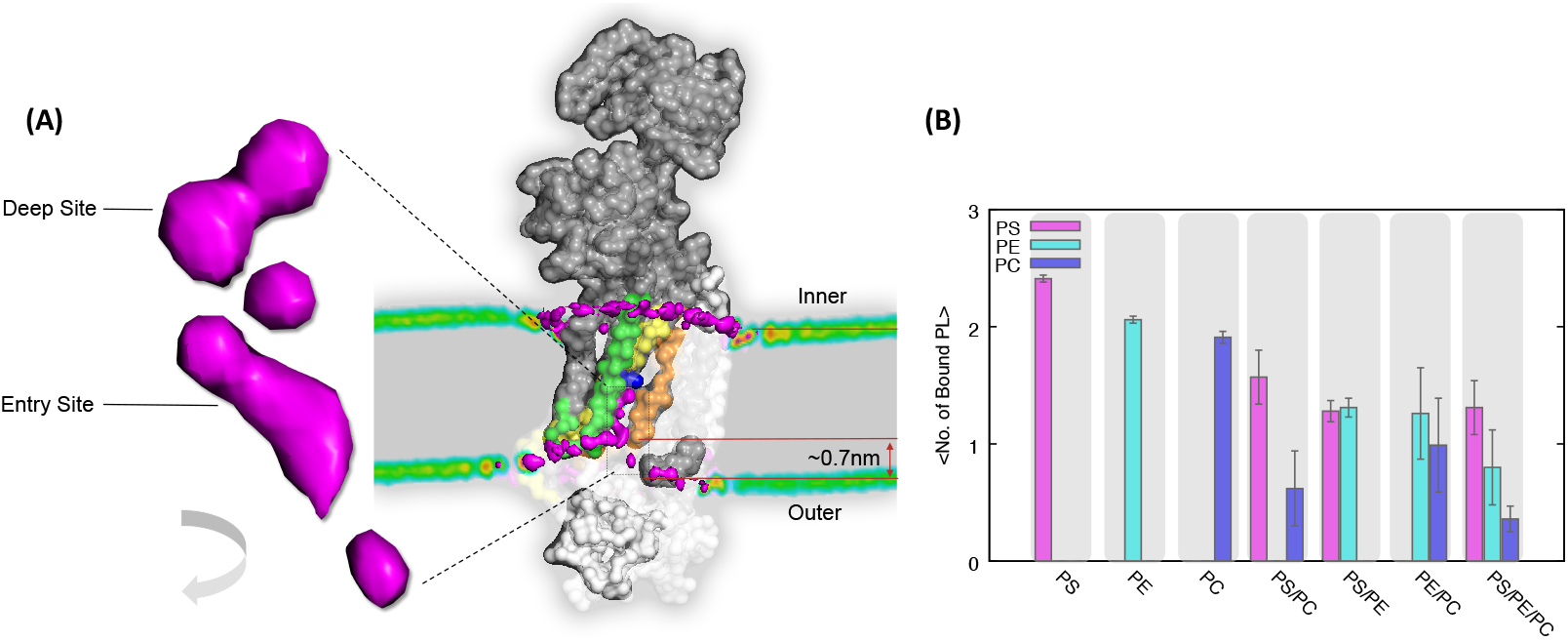
Lipid binding sites and local membrane deformation. (A) The average density map of the phosphate head group of bound lipids obtained from a 100 *µs* CG simulation with the PS/PE/PC mixed outer leaflet model (CG5 in Table 1). The average density maps obtained from other independent simulations and with different models are highly similar (Fig. S1). Drs2p and Cdc50p are presented by grey and white surfaces, respectively. TM2, TM4 and TM6 are colored in green, yellow and orange, respectively. The PISL motif in the middle of TM4 is blue. On the left is shown a zoom-in on the phosphate density map, which reveals a few binding sites which we broadly classify into a ‘deep site’ and an ‘entry site’ for simplification. The whole protein-membrane complex is shown in a slice view. The local membrane deformation is observed in the regions close to the groove, resulting in ∼0.7 nm thinning of the membrane bilayer. (B) The average number and types of lipids in the groove obtained from CG simulations with all seven different outer leaflet models. Error bars were estimated with block analysis.

Having established that the long CG simulations enable us to observe reversible lipid binding also deeper into the groove, we proceeded to examine which and how many lipids may bind. For each of the seven different setups for the outer layer composition, we counted the average number of phosphate head groups (separately for POPS/POPE/POPC as relevant for each setup) that occupy the groove (Fig. 3B) and made the following observations: (i) All PS/PE/PC lipids can bind to the groove, similar to what we observed in atomistic models; (ii) On average about two lipids may bind at the same time; (iii) The composition of the lipids inside the groove depends on the lipid composition in the outer leaflet, but is not simply proportional to the fraction of lipids in the leaflet.

Observation (iii) suggests that there is some selectivity for which lipids bind to/in the groove. In particular, it appears that the protein groove shows selectivity for PS over in particular PC lipids, but also some preference for PS over PE. This trend of POPS>POPE>POPC is in qualitative agreement with the experimental observation that PS is the natural substrate of Drs2p, PE is a weaker substrate, and PC is not a substrate. However, observation (i) suggests that the entry gate alone is not sufficient to determine the lipid selectivity, ruling out a ‘sole entry gate’ model, and suggesting selectivity is also found elsewhere in the conformational cycle. Observation (ii) agrees with the cryo-EM structures of ATP8A1 (***Hiraizumi et al., 2019***), Neo1p (***Bai et al., 2021***), Dnf1p, Dnf2p (***He et al., 2020; Bai et al., 2020***) and ATP11C (***Nakanishi et al., 2020a,b***) which together reveal electron densities suggesting multiple sites. Below, we return to the question of selectivity in the groove using free energy calculations.

### Protein induced local deformation of membrane

In addition to observing lipid binding at the single molecule level, we also examined the collective membrane dynamics to see how the protein impacts its membrane environment. We observed a remarkable local membrane deformation both in the CG (Fig. 3A and Fig. S1) and atomistic (Fig. S3) simulations. Comparable deformations are found in all independent CG simulations and models (Fig. S2), which were all initiated from a flat lipid bilayer. The deformation is only observed on the outer leaflet side and not the inner leaflet, and only in the region close to the entry of the groove. The local deformation of the membrane near the entry site to the lipid binding groove resulted in a ∼0.7-1.0 nm thinner membrane in this region, thereby likely facilitating the entry of lipid headgroups into the groove and shortening the path for lipid transport, reminiscent of the membrane distortion observed in scramblases (***Bethel and Grabe, 2016; Kalienkova et al., 2019***).

### Lipid binding preferences by alchemical free energy calculations

In the CG simulations described above we observed extensive and reversible binding of lipids in the entry site of the binding groove, whereas we only observed limited lipid exchange in the deep site. This in turn made it difficult to quantify the preference and statistics of lipid binding to this deep site, and also any potential coupling between binding at the entry and deep site. To help overcome the sampling problem in unbiased simulations, we used non-equilibrium alchemical free energy calculations (***Crooks, 1999; Gapsys et al., 2015a; Aldeghi et al., 2019***) to estimate relative binding free energies for the three different types of lipids and water.

Considering that the binding preference of a lipid in one site may also be impacted by what occupies the neighboring site, we (for each of the two sites) performed the free energy calculations in four different situations in which the neighboring site is occupied by either POPS, POPE, POPC or by water (but not any PL). First, we focused on the relative free energy difference between binding the different PLs to the deep binding site with the entry site occupied by water. The calculations support and quantify our observation that POPS binds to the deep site stronger than both POPE and POPC by 5.0 kJ/mol and 8.4 kJ/mol, respectively, when the entry site is filled with water (Fig. 4A). Therefore the non-equilibrium thermodynamic integration method provides us with additional and quantitative information about substrate preferences in the deep binding site which was difficult to obtain from the unbiased MD simulations.

**Figure 4.**
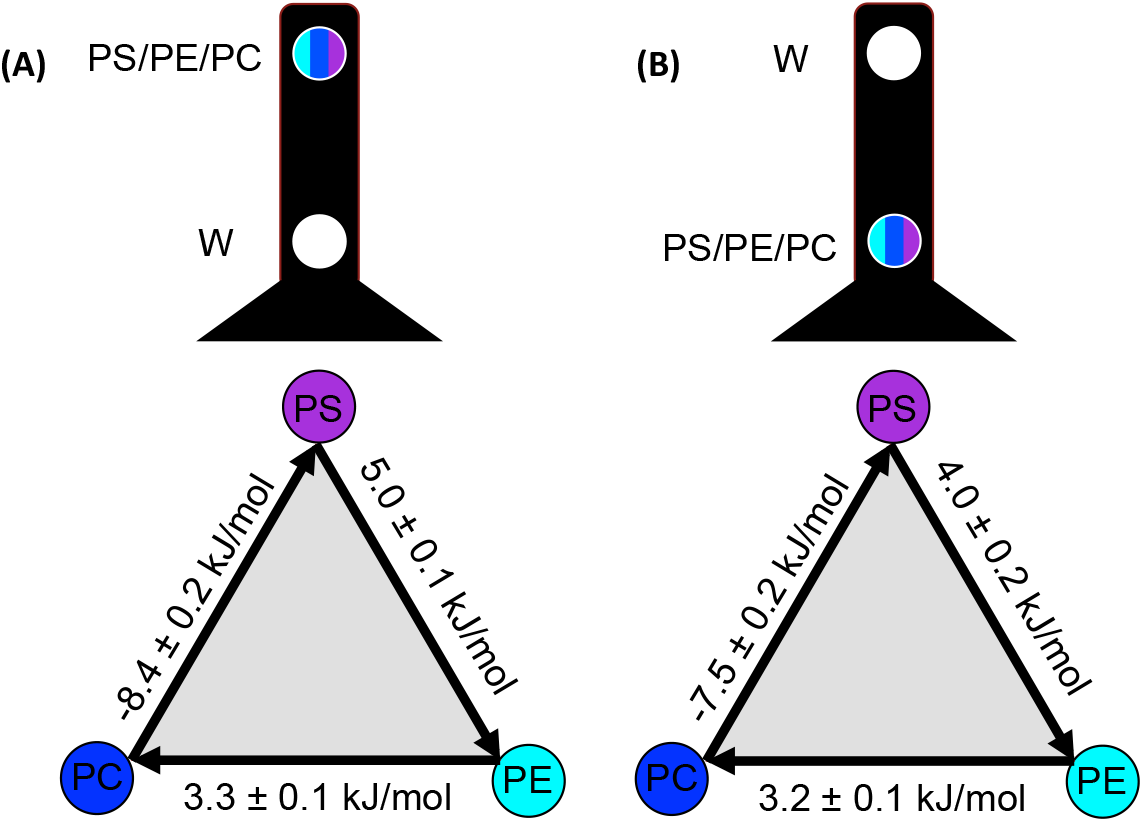
Substrate binding preferences by free energy calculations. (A) Relative binding free energy differences between POPS, POPE and POPC in the deep site in the case when the entry site is occupied by water (W). (B) Relative binding free energy differences in the entry site when the deep site is occupied by water.

To examine how the lipid selectivity in the deep site depends on the molecule bound at the entry site, we also performed the free energy calculations on the situations with either POPS/POPE/POPC occupying the entry site (Fig. S4). The results show that the magnitudes, and in a single case also the sign, of the ΔΔ*G* values depend on which molecule is bound at the neighbouring site. In all but one case, POPS binds stronger than both POPC and POPE, though the difference is greatest when the entry site is filled with water. When the entry site is filled with POPS, the situation reverses and the deep site favours (by 0.9 kJ/mol) POPE over POPS, presumably because of repulsive interactions between the charged headgroups in POPS. Nevertheless, the overall picture is that the deep site favours POPS over POPE, which in turn binds stronger than POPC. We also note that although each free energy difference was calculated independently, the thermodynamic cycles are fully consistent, so that the free energy over a full cycle sums to zero up to statistical uncertainty, supporting the notion of convergence in the calculations. We also performed free energy calculations on the entry site, confirming the preference in the order of POPS>POPE>POPC (Fig. 4B and Fig. S4B), and obtained results that were consistent with the trends we observed from the equilibrium CG simulations (Fig. 3B).

We also attempted comparable alchemical free energy calculations using atomistic simulations (Fig. S5). Despite extensive simulations (a total of 6.4 *µ*s for each PL transition), these simulations did not converge as judged for example by the fact that the free energy differences do not sum to zero over the thermodynamic cycles (sums up to ∼7-9 kJ/mol).

The binding calculations described above capture the known preference for transporting different lipids (POPS>POPE>POPC), lending support to our computational model. Nevertheless, one might still ask whether the higher affinity to POPS is an artefact of the simulation model as it has been shown to give rise to higher affinity of anionic lipids to membrane proteins (***Duncan et al., 2020***). To examine whether the affinity is simply determined by the charge, we also performed the free energy calculations on the substitution of POPS to other charged PLs, including POPG, POPA and two artificial molecules (Fig. S6). At the CG level, POPG has the same negative charge as POPS but with a less polar head group, POPA carries more negative charges than POPS but without a polar head group, while the other two artificial molecules are the same as POPS except that their CG phosphate groups carry different charges (−2e and 0e, respectively). Our calculations revealed that POPS binds to the deep site more energetically favorable than both POPG and POPA by 4.9 kJ/mol and 15.8 kJ/mol, respectively, and a POPS-like molecule with an extra negative charge also binds more weakly (Fig. S6). Thus, we suggest that even at the CG level, the determined differences in affinity and specificity are not just solely a consequence of electrostatic interactions.

### Lipid binding to the deep site is more energetically favorable than to the entry site

The alchemical free energy calculations quantified the preference of one PL relative to another at the same binding site; however they cannot tell us the relative affinity of the same lipid at the two different (entry and deep) sites. The unbiased MD simulations did also not answer this question because of the competition between different PLs and the slow PL exchange in the deep site. Thus, to examine which site in the groove is more energetically favorable for the binding of a specific PL, we used restraints to avoid the exchange of lipids between the groove and the outer leaflet. In this way, a single PL may freely explore the groove without competition with other PLs. This was achieved by performing MD simulations of the original POPS/POPE/POPC mixed outer layer model in the presence of restraints, and the results reveal that lipids favour the deep binding site (∼8.0 kJ/mol) over the entry site (Fig. S7). Thus, our results suggest that the entry site is a transient intermediate towards to the deep site, and reveal a picture that once the substrate enters the groove, it will move to the deep site driven by the free energy gradient.

### Lipid head group slides in the groove with the tails anchored in the centre of the bilayer

The recent spur of high resolution structures of P4 flippases has provided a wealth of information about the overall structure and conformational changes in this class of proteins. They have, however, only provided limited information about the interactions between the protein and the lipid substrates. X-ray diffraction and cryo-EM experiments on ATP8A1, ATP11C, Neo1p and Dnf1p/Dnf2p show density for PL head groups, but it is not possible to see the whole hydrophobic acyl chains due to their high flexibility. Our MD simulations revealed that the TM2-TM4-TM6 groove on the surface of Drs2p could accommodate one or more PL head groups whose binding locations might imply a sequential transport path from the outer layer towards the central PISL motif. Here the PL would likely trigger dephosphorylation and conformational changes leading to occlusion and subsequently opening to the inner leaflet.

The observation that only the head group of the PL is bound in the groove is consistent with the so-called ‘credit card’ model (***Pomorski and Menon, 2006; Morra et al., 2018***). The MD simulations, however, provide us with a unique opportunity to see how the lipids orient their hydrophobic tails during binding and initial transport as their head groups move towards the PISL motif. We thus used all-atom steered MD (SMD) simulations (AA5 model) to study the process of the lipid moving from the entrance of the groove to the deep site and PISL motif. Specifically, we accelerated the movement of the lipid by biasing the phosphate head group to move towards the deep site, while not directly biasing the flexible tails of the lipid molecule. Examining ten such SMD simulations we find that the ends of the lipid tails stay near the middle of the membrane bilayer while the PL head group moves up, thus changing the PL configuration from ‘vertical’ to more ‘horizontal’ (Fig. 5A). We observed a similar change of the conformation of the PL in an unbiased CG simulation (Fig. S8). Together, these results suggest that the role of the flippase is to transport the polar head group, which in itself would not easily pass the membrane, while the hydrophobic tails stay in contact with the lipid bilayer during the transport process, providing direct support of a ‘credit-card’-like model, at least for the half-reaction we examined. We also observe that the PL head group stays hydrated as it moves up the grove, suggesting that water plays an integral role in lipid transport (Fig. 5B; see also discussion further below).

**Figure 5.**
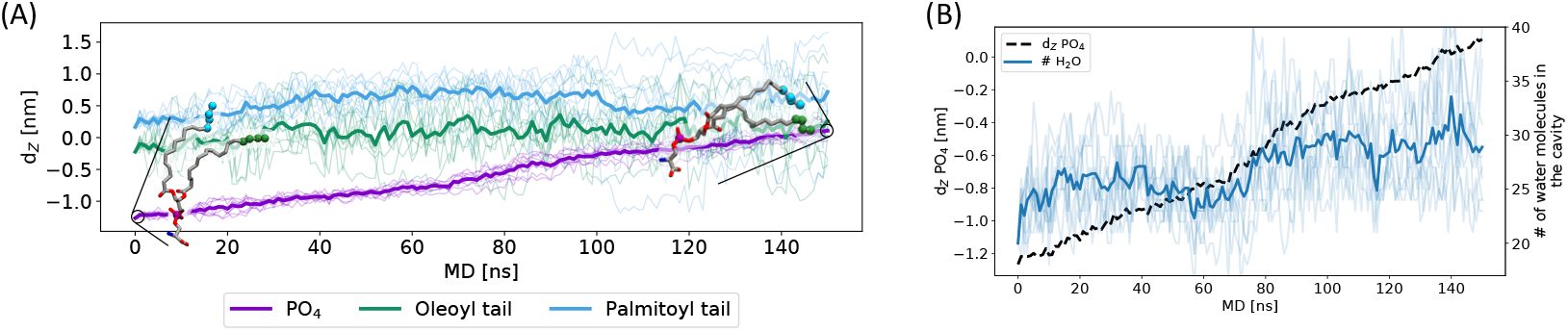
Dynamics and solvation of the lipid as it moves up in the groove. (A) Locations (along the Z axis) of the phosphate head group (purple) and ends of the two acyl chains (green and cyan) during ten 150 ns-long atomistic SMD simulations with the POPS/POPE/POPC mixed outer leaflet model (AA5). In these simulations the phosphate head group was biased to move towards the deep site during the SMD simulations while the lipid tails were left unrestrained. The thin lines show the results from each simulation while the thick lines represent the average locations. The results show how the POPS lipid performs an approximate 90-degree rotation of the whole lipid as the head group moves up the groove. (B) Number of water molecules in the cavity (right axis; dark blue line shows average while thin blue lines show each of the simulations) and phosphate location along the Z axis (left axis; black dashed line) during the SMD simulations. In both A and B, d_*z*_ = 0 corresponds to the center of the membrane while d_*z*_ = -1.1 nm corresponds to the lipid head group being in the luminal entry site.

### Two separated water-filled cavities formed within TM domain

The next question we asked was if water plays a role during the lipid binding process. To address this question, we continued to use an atomistic model (AA1) to enable a more fine-grained description of the water molecules in the groove. Starting from the cryo-EM structure of the Drs2p-Cdc50p complex with a dry TM core, we observed a substantial amount of water permeate quickly through the small gap between TM1-3 (from the inside) and the slightly bigger gap formed between TM1-TM2 loop and TM5-TM6 loop (from the outside) to fill two separated cavities in the hydrophobic TM core of Drs2p. These are formed between TM1, TM2, TM3 and TM4 near the inner layer and between TM1, TM2, TM4 and TM6 near the outer layer, respectively (Fig. 6A). These two water-filled cavities are separated by the side chain of Ile508 in the PISL motif, and we did not observe any exchange of water between the cavities, highlighting the possible important role of the PISL motif as a hydrophobic gate (***Vestergaard et al., 2014***).

**Figure 6.**
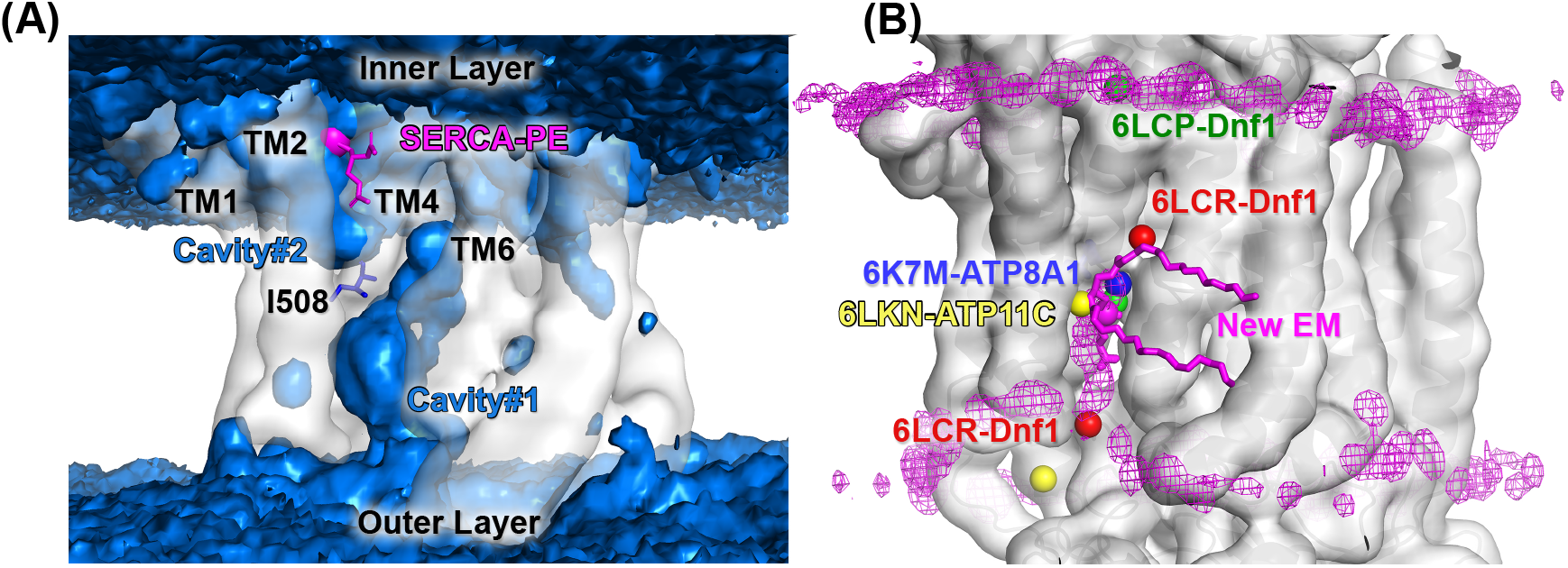
Water-filled cavities and lipid binding regions in P4 flippases. (A) Average density map of the oxygen atoms in the water molecules (blue surface) obtained from a 500 ns atomistic MD simulation with the pure POPS outer layer model (AA1 model) revealing cavities near the outer layer (overlapping with the TM2-TM4-TM6 groove) and inner layer (formed between TM1, TM2 and TM4). The Ile508 residue, shown in blue sticks, is a part of the PISL motif that separates these two water-filled cavities. (B) The average density map of the phosphate head group obtained from coarse-grained simulations with the CG5 model (magenta mesh) is superposed with the location of PL in several experimentally-determined flippase structures. These structures include our newly determined cryo-EM structure of a PL-bound E2P state of Drs2p, where we were able to model a substantial part of the POPS head group and tails (shown by sphere and sticks in magenta), as well as the PL phosphates in the E2Pi state of ATP8A1 (PDB 6K7M (***Hiraizumi et al., 2019***), blue sphere), the E2P state of ATP11C (PDB 6LKN (***Nakanishi et al., 2020a***), yellow), the E2P and E1-ATP states of Dnf1 (PDB 6LCP and 6LCR (***He et al., 2020***), green and red).

To examine more closely the functional role of the PISL motif as a seal between these two water-filled cavities, we truncated the side chain of Ile508 to generate the both Ile508Ala and Ile508Gly variants and investigated these by atomistic simulations. In simulations of both variants the two cavities merged, and we rapidly observed water exchange between them after in tens of nanoseconds (Fig. S9), demonstrating the key role of Ile508 in the gate.

A hydrated groove may be an important prerequisite for binding and transport of polar or charged lipid head groups. In the SMD simulations, we found that the number of water molecules in the cavity did not decrease as the PL head group moved up towards the PISL motif; instead we found a small increase of the number of water molecules accompanying the PL head group (Fig. 5B). This suggests a coupling between lipid binding and hydration of the groove, and may imply an important role of water on lipid movement by hydrating the polar head group. We note that in the CG MD simulations, we also observed a water-filled cavity in the outer leaflet side although the number of water molecules was less than that observed in the atomistic simulations, even after correcting for the fact that one CG water molecule corresponds to four actual water molecules (Fig. S10). We did not observe the cavity on the inner leaflet side to be filled by water in the CG simulations, perhaps because the gap on this side is too narrow to allow the bigger CG water to enter.

### Cryo-EM structure of a PS-bound PI4P-activated E2P state

Having examined the pathway by which PLs bind to and move up the groove we next set out to analyse the structural details on how Drs2p recognizes the PL head group in the deep site. In addition to the E2P structure of Drs2p (***Timcenko et al., 2019***), which formed the starting point for the simulations described above, we recently determined structures of Drs2p in several E1 states as well as a PS-occluded 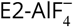 bound form (mimicking [PS]E2-Pi) (***Timcenko et al., 2021***). We here extended these studies to determine a structure of a PS-bound PI4P-activated E2P state, providing us with the possibility to examine the interactions between PL and Drs2p in the deep site (Fig. S15). We note that in contrast to our previous E2P structure (***Timcenko et al., 2019***), in which phosphorylation is mimicked by 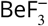, this structure was obtained through incubation with ATP and is in a phosphorylated E2P state and interestingly also has an ATP bound in the N-domain. We thus built a model of POPS-bound E2P state based on the EM map, and find that the location of the PL head group in this structure is indeed in the location predicted by our simulations (Fig. 6B). We further used this structure as starting point for a 500 ns-long unbiased all-atom simulation of this state. The simulation showed a stable complex structure, and revealed an intricate network of hydrogen bonding interactions that help stabilize the POPS head group inside the protein (Fig. 7).

**Figure 7.**
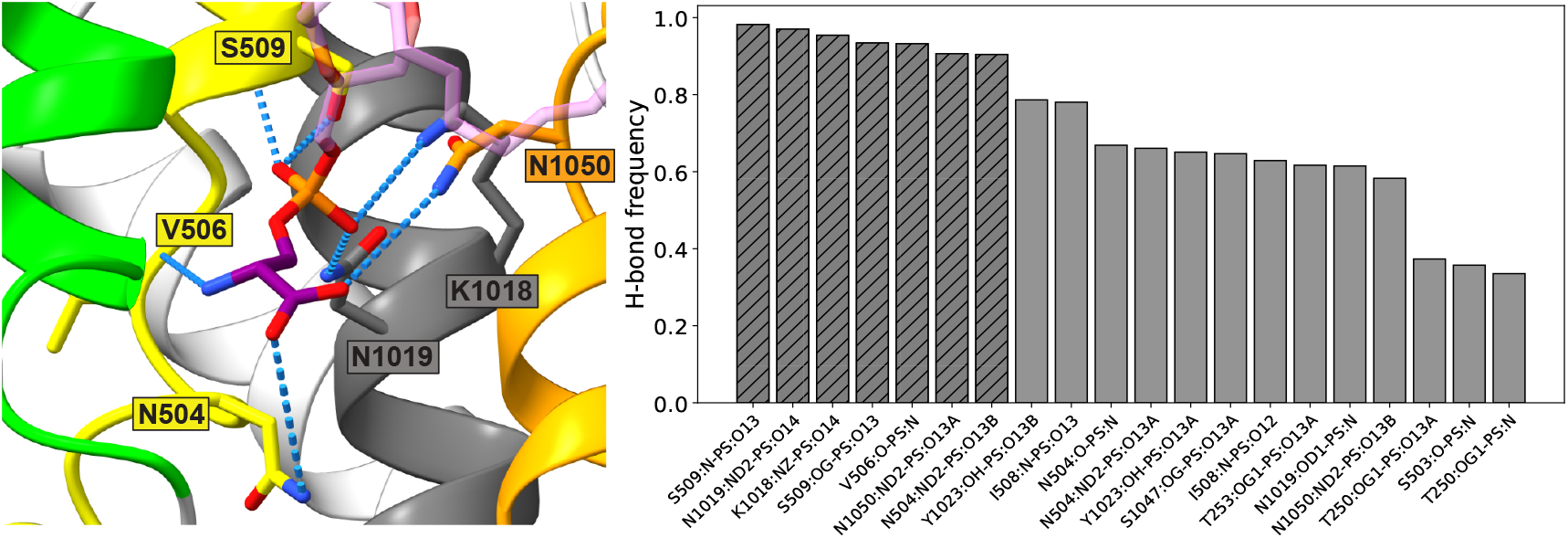
Protein-substrate interactions in a cryo-EM structure and simulations of a POPS-bound active E2P state of Drs2p. Hydrogen bonds between Drs2p and a bound POPS in the new EM structure of the active E2P state. (A) Structural representation of the seven most frequently formed hydrogen bonds (blue dashed lines) between Drs2p and the bound PS (sticks, transparent lipid tail) in the deep site. Side chains of hydrogen bond partners are represented as sticks. Residues are colored as TM4 (yellow), TM5 (grey) and TM6 (orange). (B) Fraction of time the 20 most frequent hydrogen bonds are formed between Drs2p and the bound PS molecule over 500 ns from an atomistic simulation. Hatched bars are the hydrogen bonds highlighted in panel A.

## Discussion

Biochemical and biophysical experiments have shed light on how the relatively large and amphiphilic PL substrates are recognized and transported unidirectionally by flippases, members of the P4 P-type ATPase family. The mechanism of recognition and transport of substrates is understood in exquisite detail for certain ion transporters in the P2-ATPase family, such as NKA and SERCA. It is, however, much less clear for P4 flippases, although a recent series of high-resolution structures of several flippases have resulted in detailed insights into the overall architecture and basic functional cycling of this class of molecules (***Timcenko et al., 2019; Bai et al., 2019; Hiraizumi et al., 2019; Nakanishi et al., 2020a; He et al., 2020; Nakanishi et al., 2020a; Bai et al., 2020; Timcenko et al., 2021***). The EM structure of Drs2p in a PI4P-activated E2P state reveals an empty outward-open groove on the surface of Drs2p bordered by TM2, TM4 and TM6 which was speculated to serve as a pathway for lipid entry (***Timcenko et al., 2019***). The activated E2P state of Drs2p represents a functional state with its binding pocket open to the outer leaflet, so is assumed to be ready to bind the lipid substrate from the outer leaflet of the membrane as an initial step for the substrate transport. The structure, however, did not capture the lipid substrates, the membrane nor the water molecules, leaving room for further studies using complementary tools, such as molecular simulations to understand these important details.

We have here used the outward-open structure as a starting point for providing insight into the first part of the transport cycle, examining the following questions: (i) where do the lipids bind to the flippase?, (ii) how do the lipids move?, (iii) which part of the functional cycle gives rise to substrate selectivity?, (iv) does water play a role in lipid binding or transport, and (v) how is the membrane influenced by the flippase structure, and could membrane deformations play a role in function?

In line with an earlier proposal based on the cryo-EM structure of Drs2p (***Timcenko et al., 2019***) and the reported substrate-bound structures of ATP8A1 (***Hiraizumi et al., 2019***), ATP11C (***Nakanishi et al., 2020a,b***), Dnf1 (***He et al., 2020; Bai et al., 2020***) and Dnf2 (***Bai et al., 2020***), we observed that substrate lipids enter to the TM2-TM4-TM6 groove. The groove contains multiple potential binding sites which we broadly classified into an entry and deep site. We find that the location of the deep site overlaps with the density maps of the phosphorus head group of the lipids observed in our new EM structure of the E2P state of Drs2p as well as previous EM structures of the E2P_*i*_ state of ATP8A1 (***Hiraizumi et al., 2019***), the ‘E2-site1’ in the EM structure of the E2P state of Dnf1 (***He et al., 2020***) and the crystal structure of the E2P state of ATP11C (***Nakanishi et al., 2020a,b***), while the location of the entry site overlaps with the so-called ‘E1-site2’ in the E1 state of Dnf1 (***He et al., 2020***) (Fig. 6C). Note that the site close to TM3-4 loop in the crystal structure of ATP11C and the ‘E1-site1’ in the EM structure of Dnf1 were not sampled in our coarse-grained MD simulations likely because that the ATP11C structure was captured in a different lipid environment used for crystallisation while the latter site was captured in a different state. We find that all three types of lipids we examined (POPS, POPE, POPC) bind more strongly, by about 8 kJ/mol, to the deep site than the entry site, indicating that the entry site is a transiently formed intermediate site before moving to the deep site along the transport pathway. In addition, the calculated density map around the groove also suggests at least two possible routes to enter to the groove: one uses a binding site near the TM1-TM2 loop, and another, apparently more favourable, initiating from the cavity formed between the tips of TM2, TM6 and TM9.

During the binding to the deep site, the lipid slides its head group along the groove, but keeps its acyl chains projected out into the membrane bilayer, without significant lateral movement. The lipid tails remain ‘anchored’ in the middle of the membrane, so that the whole lipid rotates approximately 90 degrees as the head group moves from the outer leaflet to the deep site. In other words, during the first half of the transport process, the substrate does not need to move its hydrophobic tails much, in line with the expectation that the movement of the hydrophilic headgroup is the most energetically unfavourable step. We speculate that for the subsequent second half of the transport event, the flippase uses the same ‘lipid tail anchor’ mechanism to completely flip the substrate to the inner leaflet.

Both our unbiased simulations as well as the alchemical free energy calculations suggest a preference for binding PL substrates in overall accordance with the known substrate specificity for full transport (POPS>POPE>POPC). We, however, also find that the very poorly transported POPC can also bind to the groove. The alchemical free energy calculations suggest the free energy of PS binding is ∼3-5 kJ/mol lower than that of PE, and ∼6-8 kJ/mol lower than PC although the relative free energy differences to a certain degree depend on what molecule occupies the neighboring site. The free energy difference we found for binding of POPS vs POPE to the deep site (5 kJ/mol) corresponds to a ∼ 7-fold stronger binding of POPS. Unfortunately, the transport rates of individual lipids by Drs2p have not been determined, making it difficult to compare these numbers directly to experiments. Nevertheless, the overall order of selectivity follows that obtained from ATPase assays (***Azouaoui et al., 2014***). We suspect, however, that the selectivity obtained from binding alone is smaller than the difference in overall transport rates of PS vs PE lipids, which would suggest that some but not all of the substrate selectivity can be explained by the difference in binding to the protein in the outward open state. In that case, the remaining part of the selectivity could e.g. come from a difference in the ability to trigger dephosphorylation and eventually transition into an inward open E1 state (***Baldridge and Graham, 2013; Vestergaard et al., 2014***). Indeed, we find that PL head groups make a number of specific hydrogen-bonding interactions in the deep site, and we speculate that these are important for triggering the conformational changes.

We observed rapid water entry into two distinct cavities in our all-atom simulations, which were initiated from the cryo-EM structure with these cavities empty. We must assume that these cavities are also filled with water in the cryo-EM experiments, only not visible, so that this filling represents the experimental structures. The two water-filled cavities are located on opposite sides of the membrane and are separated by the highly conserved PISL motif, which was also previously proposed to serve as a gate to control the lipid selectivity (***Vestergaard et al., 2014***). However, given the structural differences between Drs2p and the SERCA derived homology model, the location and geometry of these water cavities are different. Importantly, during our simulations, these two cavities were strictly isolated without water exchange between them, but only to the intra- and extracellular environment, respectively, supporting the functional role of the PISL motif as a hydrophobic gate.

We then used biased, atomistic MD simulations to accelerate the lipid movement up the groove, and observed that the head group of the lipid interacts with a substantial number of water molecules during the binding process, implying the important role of water during the lipid transport. Such water-filled cavities have also previously been observed in MD simulations of a homology model of the mammalian flippase ATP8A2 (***Vestergaard et al., 2014***) and of SERCA (***Saleh et al., 2019***). Therefore it would be interesting to investigate if the presence of water-filled cavities could be a general feature in the whole family of P-type ATPases. Interestingly, some crystal structures of SERCA (PDB ID 2AGV, 3AR4, 3W5C and 3W5D) (***Obara et al., 2005; Toyoshima et al., 2011, 2013; Drachmann et al., 2014***) captured a PE molecule bound specifically in the same location that overlaps with the water-filled cavity on the inner cytosolic layer (Fig. S11). Moreover, the Drs2p-specific residue Phe511, which is four residues from the Pro507 in the PISL motif (thus also known as the ‘proline + 4’ residue), extends its side chain to this cytosolic cavity, and has been confirmed to be involved in PS selection of Drs2p and assumed to serve as the ‘exit gate’ as proposed in the ‘two-gate’ model (***Baldridge and Graham, 2012***). Thus, P4 flippases may have evolved a capacity to use this cavity to select lipids using the ‘exit gate’ residues and drive out the substrates to the inner leaflet in the subsequent E2→E1 transition.

Finally, we also observed that the protein induces a local deformation of the membrane in the outer layer around the entry point of the lipid-transport pathway. This deformation leads to a ∼0.7-1.0 nm thinning of the membrane, which we suggest might facilitate the movement of hydrophilic head groups across the membrane. A local membrane deformation has been observed in the ATP-independent bidirectional lipid scramblases both computationally and experimentally (***Bethel and Grabe, 2016; Jiang et al., 2017; Kalienkova et al., 2019***). The local deformation may facilitate the initial binding of the substrate and the hydration of the groove may further help the movement of the head group of the substrate to a deeper location which serves as a start point for accomplishing the second half of the transport cycle.

We do note some limitations in our study. First, simulations are limited by the accuracy of the force fields used and eventually need experimental testing. Our simulations may, for example, be used to guide further mutational analyses or experiments that probe the role of water in lipid transport. Second, simulations are also limited by the accessible time scales and in their ability to capture changes in ‘chemistry’ such as a change in phosphorylation or protonation. For these reasons, we focused our study on the first part of the transport process, where a PL is taken up from the outer leaflet and the head group moves towards the centre of the bilayer, next to the PISL motif. Our results suggest that substrate binding to the deep site in the groove explains some, but not all, of the substrate selectivity, and we suggest that lipid-induced conformational change of the flippase also contributes to the lipid selectivity. More computationally-demanding atomistic simulations, possibly aided by enhanced sampling techniques, are likely needed to provide more detailed insight into the coupling between protein conformational change and lipid transport during the following steps, such as dephosphorylation and the E2→E1 transitions.

Taken together, our MD simulations and EM structure provide molecular details on lipid binding, membrane deformation and groove hydration, deepen the understanding of the dynamic transport processes otherwise inaccessible from snapshot single-particle cryo-EM and X-ray crystal structures and quantify the substrate preference on the binding sites. Our results support the notion that the groove outlined by the transmembrane segments 2, 4 and 6 serves as the first half pathway for lipid translocation, and identified the presence of multiple lipid binding sites in the groove. The results suggest the substrate preference is not determined by a single site but likely by the whole pathway, including the interactions that trigger occlusion by the binding coordination at the deep site and lead to dephosphorylation. Future studies are needed to determine and validate these interactions, and likely need an atomistic view of the binding of PL to the deep site and coupling to the dephosphorylation reaction.

## Materials and methods

### System preparation

We used the EM structure of the PI4P-activated E2P state (PDB code 6ROJ) to model the E2P state of the Drs2p-Cdc50p complex. The aspartate beryllium trifluoride in the cryo-EM structure was modeled as a double negatively-charged phosphorylated Asp. The Mg^2+^ ion, the PI4P molecule, and two disulfide bonds in Cdc50p were kept in the model. The terminal segments missing in the cryo-EM structure were considered as flexible tails, and not included in our model to save computational cost. On the basis of the predictions by PROPKA (***Olsson et al., 2011***), Glu550, Glu655, Glu798, Glu1042 in Drs2p were assigned to be protonated and Lys283 in Cdc50p was deprotonated.

### Atomistic simulations

We used the CHARMM36m force field (***Huang et al., 2016***) for the protein. Force field parameters for PI4P were generated using the CHARMM General Force Field (CGenFF) (***Vanommeslaeghe et al., 2009***) and the CHARMM-GUI web interface (***Qi et al., 2015***). Force field parameters for the phosphorylated Asp560 were as previously described (***Das et al., 2017; Saleh et al., 2019***).

POPS (1-palmitoyl-2-oleoyl-sn-glycero-3-phosphoserine), POPE (1-palmitoyl-2-oleoyl-sn-glycero-3-phosphoethanolamine), and POPC (1-Palmitoyl-2-oleoyl-sn-glycero-3-phosphocholine) were used as PS, PE and PC, lipids respectively. The Drs2p-Cdc50p complex was embedded into a series of asymmetric lipid bilayers with 67%POPE:33%POPS in the cytosolic leaflet and different compositions of mixed POPC, POPE and POPS in the luminal leaflet (Table 1). To mimic a yeast-like membrane, POPA (1-palmitoyl-2-oleoyl phosphatidic acid), POPI (1-palmitoyl-2-oleoyl phosphatidylinositol), POPG (1-palmitoyl-2-oleoyl phosphatidylglycerol) and ergosterol (ERG) were mixed together with POPS, POPE and POPC (AA6 model). The systems were solvated with a 0.1 M NaCl aqueous solution, resulting in ∼340,000 atoms in total. The dimensions of the equilibrated box were 13.8 nm, 13.8 nm and 17.1 nm in the x, y and z dimension, respectively (Fig. S12).

The temperature was kept constant at 310 K using the Nose-Hoover thermostat with a 1 ps coupling constant, and the pressure at 1.0 bar using the Parrinello-Rahman barostat with a 5 ps time coupling constant. We used a cutoff of 1.2 nm for the van der Waals interactions using a switching function starting at 1.0 nm. The cutoff for the short-range electrostatic interactions was at 1.2 nm and the long-range electrostatic interactions were calculated by the means of the particle mesh Ewald decomposition algorithm with a 0.12 nm mesh spacing. A reciprocal grid of 120× 120 × 144 cells was used with 4th order B-spline interpolation.

### Coarse-grained modeling and simulations

The atomic coordinates of the Drs2p-Cdc50p complex in E2P state were converted to the CG Martini representation by the modified martinize.py script (version 2.6) (***De Jong et al., 2013***) obtained from the MARTINI website (http://www.cgmartini.nl/). The ElNeDyn elastic-network approach was employed to restrain the protein secondary and tertiary structure, using a force constant of 500*kJ* /*mol*/*nm*^*2*^and a cut off of 0.9 nm. Position restraints were further applied to the backbone beads to keep the system in the activated E2P state and also facilitate the analysis on the lipid binding. The CG structure was embedded in a series of asymmetric mixed lipid membranes (Table 1) and solvated with MARTINI water using insane.py in a 12.5×12.5×18*nm*^3^ box (Fig. S12). To prevent unwanted freezing, 10% of the MARTINI water particles were replaced by the ‘antifreeze’ water particles. The systems were neutralized by adding NaCl at a concentration of 0.15 M.

We used the MARTINI force field version 2.2 for all CG simulations. Standard MD parameters (http://cgmartini.nl/images/parameters/exampleMDP/martini_v2.x_new-rf.mdp) with minor adjustments were used to set up the simulations. Briefly, the simulations were performed in the semi-isotropic NPT ensemble using the v-rescaling thermostat to maintain temperature at 310 K and the Parrinello-Rahman barostat to control the pressure at 1 bar with a compressibility of 3*×*10^−4^*bar*^−1^ and a coupling constant of 12 ps. Electrostatic interactions were treated using a reaction field with a dielectric constant of 15. The non-bonded interactions were cut-off at 1.1 nm. The simulations were performed using a 20 fs integration time step. The Verlet neighbor search algorithm was used to allow for GPU-accelerated simulations.

All atomistic and CG MD simulations were performed using Gromacs (***Abraham et al., 2015***) 2018 or 2019 if not specially specified. Trajectories were saved and analyzed every 1 ns.

### Alchemical free energy calculations

We used non-equilibrium alchemical free energy calculations (***Gapsys et al., 2015a; Aldeghi et al., 2019***) to determine the relative free energy difference between different substrates in a single binding site. To avoid problems arising when changing the total charge, we combined the two branches of the thermodynamic cycle into a single transition by performing *PLA*_*bilayer*_ → *P LB*_*bilayer*_ and *PLB*_*site*_ → *P LA*_*site*_ transitions at the same time (Fig. S13). By using this so-called ‘double-system/single-box’ setup (***Aldeghi et al., 2019***), we obtained ΔΔ*G* in a single set of calculations without the need of separate calculations of ΔΔ*G*_3_ and ΔΔ*G*_4_ (Fig. S13). In these calculations, we thus mutated one lipid in the outer layer (chosen to be at least 5 nm away from the protein), while the opposite transformation was performed for a lipid bound in the substrate-binding groove.

We used a single topology approach to define a hybrid topology that contains both end states for the alchemical transformations. First, we extracted the equilibrium ensembles of the CG system for the corresponding end states from the 100 *µs* long MD simulations. We then performed 100 4-ns long fast-growth transition simulations starting from 100 equally spaced frames from the equilibrium ensembles. We performed the procedure twice with a different reference lipid in the outer layer to examine the dependence on the local membrane environment, so resulting in 400 simulations (100 transitions × 2 directions × 2 reference lipids) for each ΔΔ*G* estimation. The work performed by the system was then calculated by numerical integration of the *∂H*/*∂A* curves for each transition. Finally, the free energy difference between the two end states was estimated from the overlap of the two accumulated work distributions for both directions using the Bennett Acceptance Ratio (BAR) estimator (***Shirts et al., 2003***). Uncertainties were estimated as the standard errors of the BAR estimates by bootstrap analysis on all 200 related non-equilibrium trajectories using the pmx tool (***Gapsys et al., 2015b***). All alchemical simulations were carried out using Gromacs 2019.5.

### Steered MD simulations

As the starting point for a set of ten 150 ns-long steered MD simulations we used an equilibrated system from the AA5 simulations (Table 1) with a POPS in close proximity to the entry site on the outer leaflet of the membrane. Simulation parameters were were as described above. We used GROMACS 2019.6 with PLUMED v2.5 (***Bonomi et al., 2009; Tribello et al., 2014***) for the simulations. The bias was implemented with MOVINGRESTRAINT using the z-component of the distance between the phosphate of a POPS in the luminal entry site and the center of the membrane (C*α* of ASN504 and LEU505) to steer the lipid to the centre of the membrane with *1e* = 1000*kJ* /*mol*/*nm*^2^. The number of water molecules in the cavity was counted using VMD v1.9.4a43 (***Humphrey et al., 1996***). Briefly, the cavity was represented by two spheres around the center of mass between the CA atoms of THR253, SER509, THR1045 (COM1) and THR249, ASN504, SER 1043 (COM2), respectively. The sphere radii were 1 nm and 0.8 nm for COM1 and COM2, respectively. The water molecules within the two spheres were counted every 1 ns.

### Cryo-EM structure of the PS bound E2P state of Drs2p-Cdc50p

#### Grid preparation and data collection

Drs2p-Cdc50p was purified as previously described (***Timcenko et al., 2019***). To capture a PS-bound phosphorylated E2P state, we incubated 0.6 mg/mL Drs2p-Cdc50p in 0.03 mg/mL lauryl maltose neopentyl glycol (LMNG) with 0.1 mg/mL 16:0-18:1 PS (POPS) in LMNG (resulting in a final LMNG concentration of 0.09 mg/mL) for 1 hr.

C-flat Holey Carbon grids, CF-1.2/1.3-4C (Protochips) were glow discharged for 45 s at 15 mA just prior to use. Right before 3 *µ*L sample was added to the grid, 0.6*µ*L of 20mM adenosine triphosphate (ATP) was added. The grids were vitrified on a Vitrobot IV (Thermo Fisher Scientific) at 4 °C and 100% humidity.

We acquired a total of 9837 movies from two separate grids on a Titan Krios G3i (EMBION Danish National cryo-EM Facility — Aarhus node) with an X-FEG operated at 300 kV on a Gatan K3 camera with a Bioquantum energy filter at a slit width of 20 eV. We collected 1.5 s exposures in super-resolution mode using aberration free image shift data collection (AFIS) in EPU (Thermo Fisher Scientific) at a physical pixel size of 0.66 Å/pixel (magnification of 130,000x) with a total electron dose of 60 e^−^/Å^2^.

#### Data processing

We performed data processing in cryoSPARC v2 (***Punjani et al., 2017***) (and v3 for later steps). Patch Motion Correction and Patch CTF Estimation were performed, after which 991 micrographs were rejected based on a poor CTF fit, thick ice, or too much movement, leaving 8846 micrographs for further processing. Initially the blob picker, using a cylindrical reference was used to pick 32,802 particles from 300 micrographs. These were used for generating picking templates from 2D classification, which were used to pick 691,521 particles from 3464 micrographs (first grid) and 2,337,881 particles from 5382 micrographs (second grid). The good 2D classes from the initial small stack were used to generate an *ab inito* 3D reference, while a poor class was used to generate a junk class. Particles were extracted in a 400-pixel box and downsampled to a 256-pixel box at 1.0312 Å/pixel. The two stacks were subjected separately to multiple rounds of heterogeneous 3D refinement with one good reference and multiple junk references. The 668,160 protein-like picks were combined and subjected to one round of *ab initio* 3D refinement, and the 531,985 particles were refined into a consensus volume with reported resolution of 2.9 Å by non-uniform (NU) refinement (***Punjani et al., 2020***).

We subjected the remaining particles to two rounds of 3D variability analysis (3DVA) (***Punjani and Fleet, 2021***), first to select for open E2P states (205,221 particles), and secondly to select for particles with PS-bound in the transport pathway. The selected 162,243 particles were subjected to two rounds of *ab initio* 3D refinement, before the final stack of 91,415 particles was refined to 3.1 Å using NU-refinement. The final density and map-to-model fit can be seen in Figs. S14 and S15. Figures were prepared using PyMOL and UCSF Chimera (***Pettersen et al., 2004***).

### Model building and MD simulation of the POPS-bound E2P state

We used the structure of Drs2p-Cdc50p in an activated 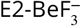 (PDB 6ROJ) as starting point for building a model of the POPS-bound E2P state using molecular dynamics flexible fitting with secondary structure, cis-peptide and chirality restraints to prevent overfitting (***Trabuco et al., 2008***). These simulations were performed using an implicit solvent with the scaling factor of the map potential *g* = 0.3. The model was refined by manual adjustment in PyMol and energy minimization in Gromacs. The refined POPS-bound structure was subsequently used to perform a 500 ns-long all-atom MD simulation using the same settings as in the simulations above.

For deposition to the Protein Data Bank (PDB), the structure was subjected to PHENIX Real Space Refine (***Afonine et al., 2018***) and manually inspected in COOT (***Emsley et al., 2010***). Validation was performed by MolProbidity (***Chen et al., 2010***) in PHENIX (***Adams et al., 2010***) and relevant metrics can be found in Table S1.

### Simulation analysis

Protein structures were visualized with PyMOL (***Schrödinger, LLC, 2015***) and analysis of simulation data was conducted using VMD (***Humphrey et al., 1996***) and GROMACS tools (***Abraham et al., 2015***), and in-house scripts. Averaged density maps of the lipid phosphate group were generated using GROMAPs (***Briones et al., 2019***) and *g*_*lomepro* (***Gapsys et al., 2013***). The interactions between Drs2p and the bound POPS were analyzed by GetContacts scripts (https://getcontacts.github.io/). Error estimates of the average number of bound PLs were determined with block analysis (***Flyvbjerg and Petersen, 1989***).

## Data availability

The cryo-EM map is deposited in The Electron Microscopy Data Bank (EMDB) with accession code EMDB-13353, and the structure is available in the Protein Data Bank with PDB ID 7PEM. Input files, simulations and analysis scripts are available at https://github.com/KULL-Centre/papers/tree/main/2020/flippase-wang-et-al-2020.

## Acknowledgments

We thank Long Li and Kazuhiro Abe for providing the coordinates of Dnf1 and ATP11C. Y.W., J.A.L., M.T., F.K. P.N. and K.L.-L. were supported by the BRAINSTRUC structural biology initiative from the Lundbeck Foundation (R155-2015-2666). Y.W. was also supported by the HPC-Europa3 Transnational Access programme and start-up funds from Zhejiang University. V.G. was supported by the BioExcel CoE (http://www.bioexcel.eu), a project funded by the European Union (Contract H2020-INFRAEDI-02-2018-823830). We acknowledge access to computational resources from the Danish National Supercomputer for Life Sciences (Computerome), the ROBUST Resource for Biomolecular Simulations (supported by the Novo Nordisk Foundation) and the HPC-Europa3 HLRS HPC center.

## References

Abraham MJ, Murtola T, Schulz R, Páll S, Smith JC, Hess B, Lindahl E. GROMACS: High performance molecular simulations through multi-level parallelism from laptops to supercomputers. SoftwareX. 2015; 1:19–25.

Adams PD, Afonine PV, Bunkóczi G, Chen VB, Davis IW, Echols N, Headd JJ, Hung LW, Kapral GJ, Grosse-Kunstleve RW, et al. PHENIX: a comprehensive Python-based system for macromolecular structure solution. Acta Crystallographica Section D: Biological Crystallography. 2010; 66(2):213–221.

Afonine PV, Poon BK, Read RJ, Sobolev OV, Terwilliger TC, Urzhumtsev A, Adams PD. Real-space refinement in PHENIX for cryo-EM and crystallography. Acta Crystallographica Section D: Structural Biology. 2018; 74(6):531–544.

Albers RW. Biochemical Aspects of Active Transport. Annual Review of Biochemistry. 1967 jun; 36(1):727–756.

Aldeghi M, de Groot BL, Gapsys V. Accurate Calculation of Free Energy Changes upon Amino Acid Mutation. In: Methods in Molecular Biology, vol. 1851 Humana Press Inc.; 2019. p. 19–47.

Andersen JP, Vestergaard AL, Mikkelsen SA, Mogensen LS, Chalat M, Molday RS. P4-ATPases as Phospholipid Flippases—Structure, Function, and Enigmas. Frontiers in Physiology. 2016 jul; 7:275.

Ansell TB, Curran L, Horrell MR, Pipatpolkai T, Letham SC, Song W, Siebold C, Stansfeld PJ, Sansom MS, Corey RA. Relative Affinities of Protein-Cholesterol Interactions from Equilibrium Molecular Dynamics Simulations. bioRxiv. 2021;.

Azouaoui H, Montigny C, Ash MR, Fijalkowski F, Jacquot A, Grønberg C, López-Marqués RL, Palmgren MG, Garrigos M, Le Maire M, et al. A high-yield co-expression system for the purification of an intact Drs2p-Cdc50p lipid flippase complex, critically dependent on and stabilized by phosphatidylinositol-4-phosphate. PLoS One. 2014; 9(11).

Bai L, Jain BK, You Q, Duan HD, Graham TR, Li H. Structural basis of the P4B ATPase lipid flippase activity. bioRxiv. 2021;.

Bai L, Kovach A, You Q, Hsu HC, Zhao G, Li H. Autoinhibition and activation mechanisms of the eukaryotic lipid flippase Drs2p-Cdc50p. Nature communications. 2019; 10(1):1–10.

Bai L, You Q, Jain BK, Duan HD, Kovach A, Graham TR, Li H. Transport mechanism of P4 ATPase phosphatidyl-choline flippases. eLife. 2020 dec; 9.

Baldridge RD, Graham TR. Identification of residues defining phospholipid flippase substrate specificity of type IV P-type ATPases. Proceedings of the National Academy of Sciences. 2012; 109(6):E290–E298.

Baldridge RD, Graham TR. Two-gate mechanism for phospholipid selection and transport by type IV P-type ATPases. Proceedings of the National Academy of Sciences. 2013; 110(5):E358–E367.

Bethel NP, Grabe M. Atomistic insight into lipid translocation by a TMEM16 scramblase. Proceedings of the National Academy of Sciences of the United States of America. 2016; 113(49):14049–14054.

Bonomi M, Branduardi D, Bussi G, Camilloni C, Provasi D, Raiteri P, Donadio D, Marinelli F, Pietrucci F, Broglia RA, Parrinello M. PLUMED: A portable plugin for free-energy calculations with molecular dynamics. Computer Physics Communications. 2009 oct; 180(10):1961–1972.

Briones R, Blau C, Kutzner C, de Groot BL, Aponte-Santamaría C. GROmaps: A GROMACS-Based Toolset to Analyze Density Maps Derived from Molecular Dynamics Simulations. Biophysical Journal. 2019 jan; 116(1):4–11.

Carafoli E, Calcium signaling: A tale for all seasons. National Academy of Sciences; 2002.

Chen VB, Arendall WB, Headd JJ, Keedy DA, Immormino RM, Kapral GJ, Murray LW, Richardson JS, Richardson DC. MolProbity: all-atom structure validation for macromolecular crystallography. Acta Crystallographica Section D: Biological Crystallography. 2010; 66(1):12–21.

Corradi V, Mendez-Villuendas E, Ingólfsson HI, Gu RX, Siuda I, Melo MN, Moussatova A, DeGagné LJ, Sejdiu BI, Singh G, Wassenaar TA, Delgado Magnero K, Marrink SJ, Tieleman DP. Lipid–Protein Interactions Are Unique Fingerprints for Membrane Proteins. ACS Central Science. 2018 jun; 4(6):709–717.

Corradi V, Sejdiu BI, Mesa-Galloso H, Abdizadeh H, Noskov SY, Marrink SJ, Tieleman DP. Emerging diversity in lipid–protein interactions. Chemical reviews. 2019; 119(9):5775–5848.

Crooks GE. Entropy production fluctuation theorem and the nonequilibrium work relation for free energy differences. Physical Review E - Statistical Physics, Plasmas, Fluids, and Related Interdisciplinary Topics. 1999 sep; 60(3):2721–2726.

Das A, Rui H, Nakamoto R, Roux B. Conformational Transitions and Alternating-Access Mechanism in the Sarcoplasmic Reticulum Calcium Pump. Journal of Molecular Biology. 2017 mar; 429(5):647–666.

Daum G, Tuller G, Nemec T, Hrastnik C, Balliano G, Cattel L, Milla P, Rocco F, Conzelmann A, Vionnet C, Kelly DE, Kelly S, Schweizer E, Schüller HJ, Hojad U, Greiner E, Finger K. Systematic analysis of yeast strains with possible defects in lipid metabolism. Yeast. 1999 may; 15(7):601–614.

De Jong DH, Singh G, Bennett WFD, Arnarez C, Wassenaar TA, Schäfer LV, Periole X, Tieleman DP, Marrink SJ. Improved parameters for the martini coarse-grained protein force field. Journal of Chemical Theory and Computation. 2013 jan; 9(1):687–697.

Drachmann ND, Olesen C, Møller JV, Guo Z, Nissen P, Bublitz M. Comparing crystal structures of Ca^2+^ -ATPase in the presence of different lipids. FEBS Journal. 2014 sep; 281(18):4249–4262.

Duncan AL, Corey RA, Sansom MSP. Defining how multiple lipid species interact with inward rectifier potassium (Kir2) channels. Proceedings of the National Academy of Sciences. 2020; 117(14):7803–7813.

Dyla M, Kjærgaard M, Poulsen H, Nissen P. Structure and Mechanism of P-Type ATPase Ion Pumps. Annual Review of Biochemistry. 2020 jun; 89(1):annurev–biochem–010611–112801.

Emsley P, Lohkamp B, Scott WG, Cowtan K. Features and development of Coot. Acta Crystallographica Section D: Biological Crystallography. 2010; 66(4):486–501.

Fadok VA, Voelker DR, Campbell PA, Cohen JJ, Bratton DL, Henson PM. Exposure of phosphatidylserine on the surface of apoptotic lymphocytes triggers specific recognition and removal by macrophages. Journal of immunology (Baltimore, Md : 1950). 1992 apr; 148(7):2207–16.

Falzone ME, Malvezzi M, Lee BC, Accardi A. Known structures and unknown mechanisms of TMEM16 scramblases and channels. The Journal of General Physiology. 2018 jul; 150(7):933–947.

Flyvbjerg H, Petersen HG. Error estimates on averages of correlated data. The Journal of Chemical Physics. 1989; 91(1):461–466.

Gapsys V, De Groot BL, Briones R. Computational analysis of local membrane properties. Journal of Computer-Aided Molecular Design. 2013 oct; 27(10):845–858.

Gapsys V, Michielssens S, Peters JHe, de Groot BL, Leonov H. Calculation of binding free energies. Methods in molecular biology (Clifton, NJ). 2015; 1215:173–209.

Gapsys V, Michielssens S, Seeliger D, De Groot BL. pmx: Automated protein structure and topology generation for alchemical perturbations. Journal of Computational Chemistry. 2015 feb; 36(5):348–354.

He Y, Xu J, Wu X, Li L. Structures of a P4-ATPase lipid flippase in lipid bilayers. Protein & Cell. 2020; p. 1–6.

Hiraizumi M, Yamashita K, Nishizawa T, Nureki O. Cryo-EM structures capture the transport cycle of the P4-ATPase flippase. Science (New York, NY). 2019 aug; p. eaay3353.

Huang J, Rauscher S, Nawrocki G, Ran T, Feig M, De Groot BL, Grubmüller H, MacKerell AD. CHARMM36m: An improved force field for folded and intrinsically disordered proteins. Nature Methods. 2016 dec; 14(1):71–73.

Humphrey W, Dalke A, Schulten K. VMD – Visual Molecular Dynamics. Journal of Molecular Graphics. 1996; 14:33–38.

Jensen MS, Costa SR, Duelli AS, Andersen PA, Poulsen LR, Stanchev LD, Gourdon P, Palmgren M, Pomorski TG, López-Marqués RL. Phospholipid flipping involves a central cavity in P4 ATPases. Scientific Reports. 2017; 7(1):17621.

Jiang T, Yu K, Hartzell HC, Tajkhorshid E. Lipids and ions traverse the membrane by the same physical pathway in the nhTMEM16 scramblase. eLife. 2017 sep; 6.

Kalienkova V, Mosina VC, Bryner L, Oostergetel GT, Dutzler R, Paulino C. Stepwise activation mechanism of the scramblase nhtmem16 revealed by cryoem. eLife. 2019; 8:e44364.

Kobayashi T, Menon AK. Transbilayer lipid asymmetry. Current Biology. 2018 apr; 28(8):R386–R391.

Lee S, Uchida Y, Wang J, Matsudaira T, Nakagawa T, Kishimoto T, Mukai K, Inaba T, Kobayashi T, Molday RS, et al. Transport through recycling endosomes requires EHD 1 recruitment by a phosphatidylserine translocase. The EMBO journal. 2015; 34(5):669–688.

Levental KR, Malmberg E, Symons JL, Fan YY, Chapkin RS, Ernst R, Levental I. Lipidomic and biophysical homeostasis of mammalian membranes counteracts dietary lipid perturbations to maintain cellular fitness. Nature communications. 2020; 11(1):1–13.

Lipp NF, Gautier R, Magdeleine M, Renard M, Albanèse V, Copic A, Drin G. An electrostatic switching mechanism to control the lipid transfer activity of Osh6p. Nature Communications. 2019 dec; 10(1):3926.

Lyons JA, Timcenko M, Dieudonné T, Lenoir G, Nissen P, P4-ATPases: how an old dog learnt new tricks — structure and mechanism of lipid flippases. Elsevier Ltd; 2020.

Marrink SJ, Risselada HJ, Yefimov S, Tieleman DP, De Vries AH. The MARTINI force field: Coarse grained model for biomolecular simulations. Journal of Physical Chemistry B. 2007 jul; 111(27):7812–7824.

Menon I, Huber T, Sanyal S, Banerjee S, Barré P, Canis S, Warren JD, Hwa J, Sakmar TP, Menon AK. Opsin is a phospholipid flippase. Current Biology. 2011 jan; 21(2):149–153.

Montigny C, Lyons J, Champeil P, Nissen P, Lenoir G. On the molecular mechanism of flippase-and scramblase-mediated phospholipid transport. Biochimica et Biophysica Acta (BBA)-Molecular and Cell Biology of Lipids. 2016; 1861(8):767–783.

Morra G, Razavi AM, Pandey K, Weinstein H, Menon AK, Khelashvili G. Mechanisms of lipid scrambling by the G protein-coupled receptor opsin. Structure. 2018; 26(2):356–367.

Muller MP, Jiang T, Sun C, Lihan M, Pant S, Mahinthichaichan P, Trifan A, Tajkhorshid E. Characterization of lipid–protein interactions and lipid-mediated modulation of membrane protein function through molecular simulation. Chemical reviews. 2019; 119(9):6086–6161.

Nakanishi H, Irie K, Segawa K, Hasegawa K, Fujiyoshi Y, Nagata S, Abe K. Crystal structure of a human plasma membrane phospholipid flippase. Journal of Biological Chemistry. 2020 jun; p. jbc.RA120.014144.

Nakanishi H, Nishizawa T, Segawa K, Nureki O, Fujiyoshi Y, Nagata S, Abe K. Transport Cycle of Plasma Membrane Flippase ATP11C by Cryo-EM. Cell Reports. 2020 sep; 32(13):108208.

Natarajan P, Wang J, Hua Z, Graham TR. Drs2p-coupled aminophospholipid translocase activity in yeast Golgi membranes and relationship to in vivo function. Proceedings of the National Academy of Sciences of the United States of America. 2004 jul; 101(29):10614–10619.

Obara K, Miyashita N, Xu C, Toyoshima I, Sugita Y, Inesi G, Toyoshima C. Structural role of countertransport revealed in Ca2+ pump crystal structure in the absence of Ca2+. Proceedings of the National Academy of Sciences of the United States of America. 2005 oct; 102(41):14489–14496.

Olsson MHM, SØndergaard CR, Rostkowski M, Jensen JH. PROPKA3: Consistent treatment of internal and surface residues in empirical p K a predictions. Journal of Chemical Theory and Computation. 2011 feb; 7(2):525–537.

Palmgren M, Østerberg JT, Nintemann SJ, Poulsen LR, López-Marqués RL. Evolution and a revised nomenclature of P4 ATPases, a eukaryotic family of lipid flippases. Biochimica et Biophysica Acta (BBA)-Biomembranes. 2019; 1861(6):1135–1151.

Palmgren MG, Nissen P. P-Type ATPases. Annual Review of Biophysics. 2011 jun; 40(1):243–266.

Pettersen EF, Goddard TD, Huang CC, Couch GS, Greenblatt DM, Meng EC, Ferrin TE. UCSF Chimera—a visualization system for exploratory research and analysis. Journal of computational chemistry. 2004; 25(13):1605–1612.

Pomorski T, Menon AK, Lipid flippases and their biological functions. Springer; 2006.

Post RL, Hegyvary C, Kume S. Activation by adenosine triphosphate in the phosphorylation kinetics of sodium and potassium ion transport adenosine triphosphatase. Journal of Biological Chemistry. 1972; 247(20):6530–6540.

Punjani A, Fleet DJ. 3D variability analysis: Resolving continuous flexibility and discrete heterogeneity from single particle cryo-EM. Journal of Structural Biology. 2021; 213(2):107702.

Punjani A, Rubinstein JL, Fleet DJ, Brubaker MA. cryoSPARC: algorithms for rapid unsupervised cryo-EM structure determination. Nature methods. 2017; 14(3):290–296.

Punjani A, Zhang H, Fleet DJ. Non-uniform refinement: adaptive regularization improves single-particle cryo-EM reconstruction. Nature methods. 2020; 17(12):1214–1221.

Qi Y, Ingólfsson HI, Cheng X, Lee J, Marrink SJ, Im W. CHARMM-GUI Martini Maker for Coarse-Grained Simulations with the Martini Force Field. Journal of Chemical Theory and Computation. 2015 sep; 11(9):4486–4494.

Roland BP, Graham TR. Directed evolution of a sphingomyelin flippase reveals mechanism of substrate backbone discrimination by a P4-ATPase. Proceedings of the National Academy of Sciences. 2016; 113(31):E4460–E4466.

Roland BP, Naito T, Best JT, Arnaiz-Yépez C, Takatsu H, Roger JY, Shin HW, Graham TR. Yeast and human P4-ATPases transport glycosphingolipids using conserved structural motifs. Journal of Biological Chemistry. 2019; 294(6):1794–1806.

Saleh N, Wang Y, Nissen P, Lindorff-Larsen K. Allosteric modulation of the sarcoplasmic reticulum Ca 2+ ATPase by thapsigargin via decoupling of functional motions. Physical Chemistry Chemical Physics. 2019; 21(39):21991–21995.

Schrödinger, LLC. The PyMOL Molecular Graphics System, Version 1.8; 2015.

Shirts MR, Bair E, Hooker G, Pande VS. Equilibrium free energies from nonequilibrium measurements using maximum-likelihood methods. Physical Review Letters. 2003 oct; 91(14):140601.

Takatsu H, Tanaka G, Segawa K, Suzuki J, Nagata S, Nakayama K, Shin HW. Phospholipid flippase activities and substrate specificities of human type IV P-type ATPases localized to the plasma membrane. Journal of Biological Chemistry. 2014; 289(48):33543–33556.

Timcenko M, Dieudonne T, Montigny C, Boesen T, Lyons JA, Lenoir G, Nissen P. Structural basis of substrate-independent phosphorylation in a P4-ATPase lipid flippase. Journal of Molecular Biology. 2021; p. 167062.

Timcenko M, Lyons JA, Januliene D, Ulstrup JJ, Dieudonné T, Montigny C, Ash MR, Karlsen JL, Boesen T, Kühlbrandt W, Others. Structure and autoregulation of a P4-ATPase lipid flippase. Nature. 2019; p. 1.

Toyoshima C, Iwasawa S, Ogawa H, Hirata A, Tsueda J, Inesi G. Crystal structures of the calcium pump and sarcolipin in the Mg 2+-bound E1 state. Nature. 2013 mar; 495(7440):260–264.

Toyoshima C, Kanai R, Cornelius F. First crystal structures of Na+,K+-ATPase: new light on the oldest ion pump. Structure (London, England : 1993). 2011 dec; 19(12):1732–8.

Trabuco LG, Villa E, Mitra K, Frank J, Schulten K. Flexible Fitting of Atomic Structures into Electron Microscopy Maps Using Molecular Dynamics. Structure. 2008 may; 16(5):673–683.

Tribello GA, Bonomi M, Branduardi D, Camilloni C, Bussi G. PLUMED 2: New feathers for an old bird. Computer Physics Communications. 2014 feb; 185(2):604–613.

Vanommeslaeghe K, Hatcher E, Acharya C, Kundu S, Zhong S, Shim J, Darian E, Guvench O, Lopes P, Vorobyov I, Mackerell AD. CHARMM general force field: A force field for drug-like molecules compatible with the CHARMM all-atom additive biological force fields. Journal of Computational Chemistry. 2009; p. NA–NA.

Vestergaard AL, Coleman JA, Lemmin T, Mikkelsen SA, Molday LL, Vilsen B, Molday RS, Dal Peraro M, Andersen JP. Critical roles of isoleucine-364 and adjacent residues in a hydrophobic gate control of phospholipid transport by the mammalian P4-ATPase ATP8A2. Proceedings of the National Academy of Sciences. 2014; 111(14):E1334–E1343.

