## Supporting Figures for "Substrate Transport and Specificity in a Phospholipid Flippase"

[1]Structural Biology and NMR Laboratory & Linderstrøm-Lang Centre for Protein Science, Department of Biology, University of Copenhagen, Denmark; [2]Shanghai Institute for Advanced Study, Institute of Quantitative Biology, College of Life Sciences, Zhejiang University, Hangzhou 310027, China; [3]Danish Research Institute of Translational Neuroscience - DANDRITE, Nordic EMBL Partnership for Molecular Medicine, Dept. Molecular Biology and Genetics, Aarhus University, Aarhus, Denmark; [4]Computational Biomolecular Dynamics Group, Department of Theoretical and Computational Biophysics, Max Planck Institute for Biophysical Chemistry, D-37077 Göttingen, Germany

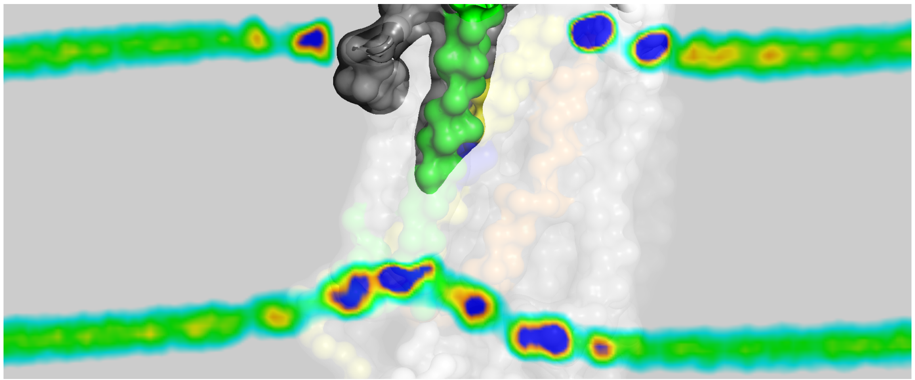

**Figure S1. Local membrane deformation induced by flippase.** The figure shows a slice view of the average density map of the phosphate head group in the phospholipid bilayer obtained from the coarse-grained PS/PE/PC mixed outer layer model (CG5) with a clipping plane (transparent and grey surface) parallel to the Z axis and close to the TM2-TM4-TM6 groove. The dark blue color indicates regions of high density.

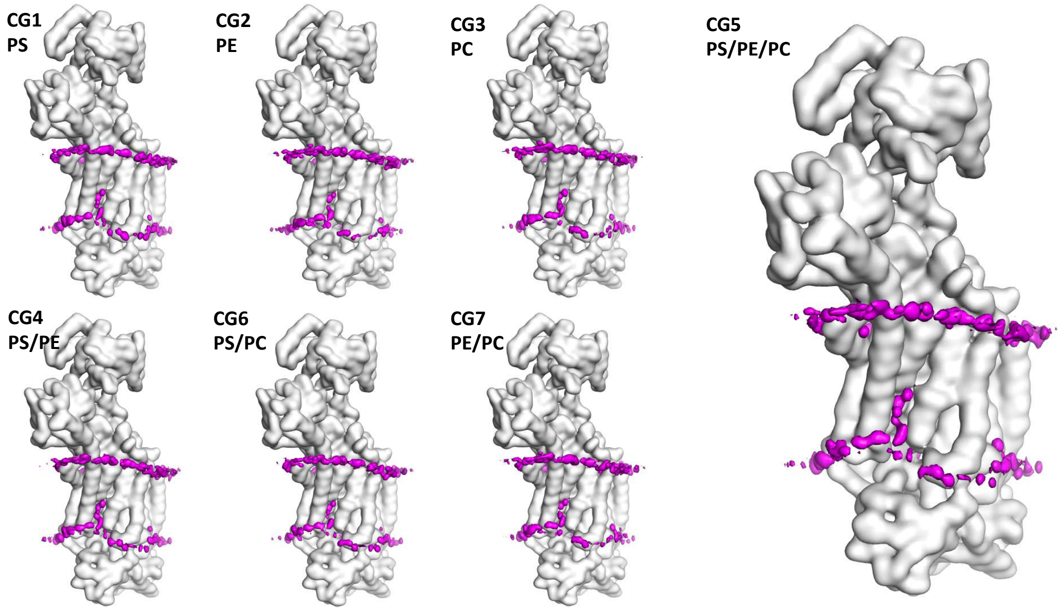


**Figure S2. The average density map of the phosphate head group of bound lipids obtained from CG simulations.** It shows the results from different mixed outer leaflet model (CG1-CG7 in Table 1 in the main text). Each map was obtained from a 100 $\mu s$ CG simulation.

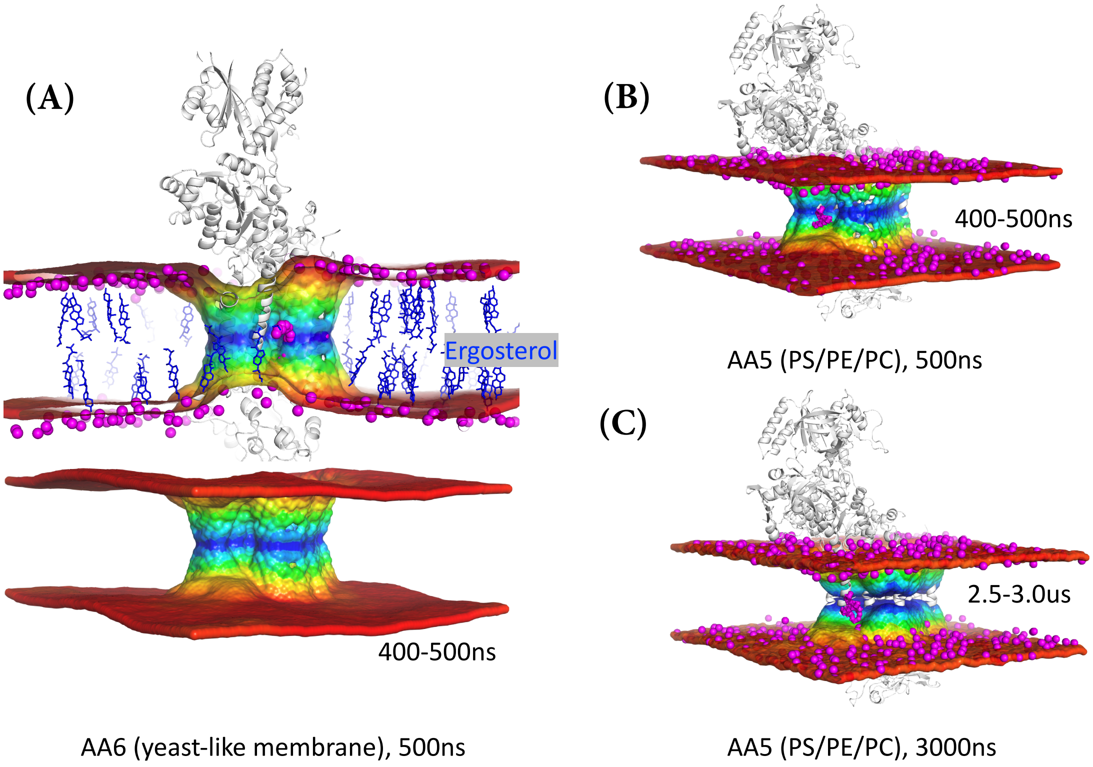


**Figure S3. Local membrane deformation observed in all-atom simulations.** (A) shows the local thickness of the membrane in an all-atom yeast-like membrane model (AA6). The last 100 ns trajectory of a 500ns MD simulation was used for the analysis. (B) shows the local thickness of the membrane in an all-atom PS/PE/PC mixed model (AA5). The last 100 ns trajectory of a 500ns MD simulation was used for the analysis. (C) shows the local thickness of the membrane in AA5 model but with longer 3$\mu s$ simulation time. The last 500 ns trajectory was used for the analysis. Membrane thickness was calculated considering the phosphorus atoms using the Local Membrane Property Analysis tool $g_{lomepro}$ and colored from red (high thickness) to blue (low thickness).

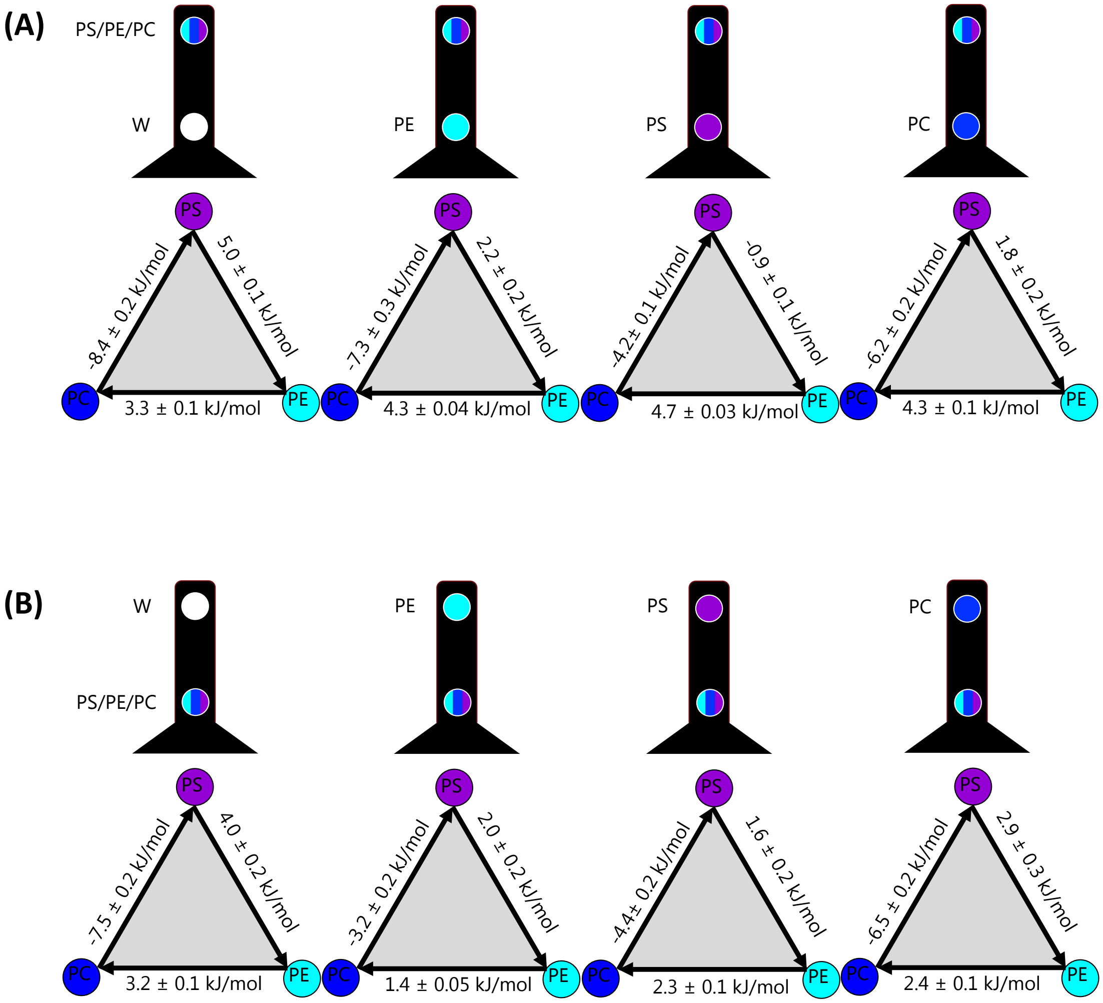


**Figure S4. Substrate binding preference by free energy calculations.** Thermodynamic free energies of binding, as also shown in Fig. 4 A: Relative free energies for binding of different lipids to the deep site, when the entry site is occupied by water, POPE, POPS or POPC, respectively. B: Relative free energies for binding of different lipids to the entry site, when the deep site is occupied by water, POPE, POPS or POPC, respectively.

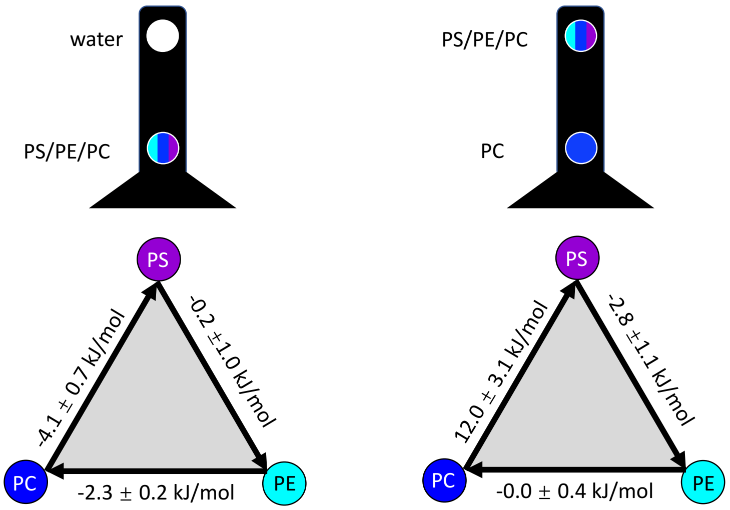


**Figure S5. Alchemical free energy calculations on both sites using the atomistic models (AA5).** We attempted to perform all-atom free energy calculations. The free energies over the full thermodynamic cycles, however, sum up to ~7-9 kJ/mol suggesting lack of convergence. This in turn means that these all-atom calculations cannot capture the subtle free energy differences and quantify the lipid specificity. The lack of convergence was probably due to the difficulty to sample the alchemical transitions between PLs, which involve the change of a substantial number of atoms, and maybe also partially be due to the challenge of equilibrating the PL-bound ensemble. This is despite performing extensive alchemical simulations (a total of 6.4 $\mu$s accumulated for each PL transition).

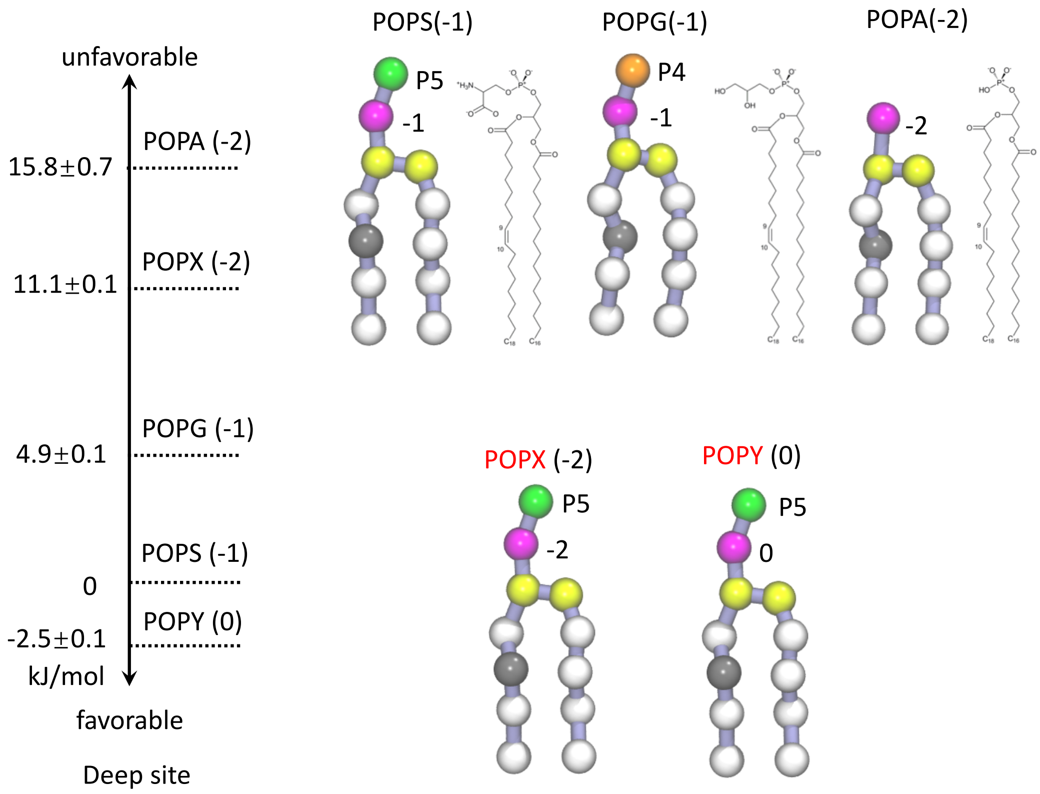


**Figure S6. Alchemical free energy calculation of PS mutations in the deep site using CG simulations (CG6).** The figure shows the free energy changes on the transitions of POPS to other charged PLs, including POPG, POPA and two artificial molecules. At the CG level, POPG has the same negative charge as POPS but with a less polar head group (with a 'P4' Martini bead), POPA carries more negative charges than POPS but without a polar head group, while the other two artificial molecules (which we called POPX and POPY) are the same as POPS except that their CG phosphate groups carry different charges (-2e and 0e, respectively).

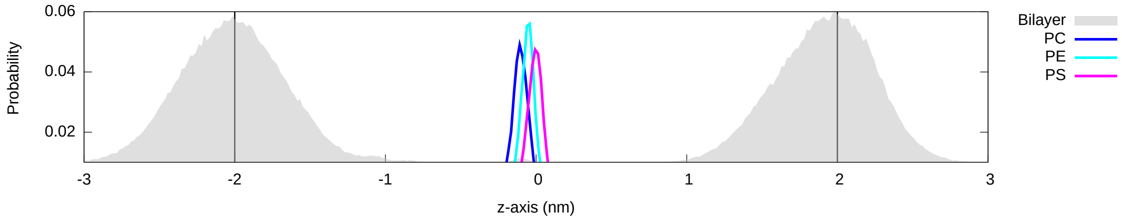


**Figure S7. The deep site binds lipids more tightly than the entry site.** To examine which site in the groove is more energetically favorable for the binding of a specific PL, we used restraints to avoid exchange of lipids between the groove and the outer leaflet. The figure shows the density probability of the PO4 head group of the bound PL along the z axis. The results were obtained from MD simulations using the restrained CG5 models in which only a single PL was allowed to freely explore the groove without competition with other PLs. As a reference, the density profile of the PO4 head group of the bilayer was shown in grey.

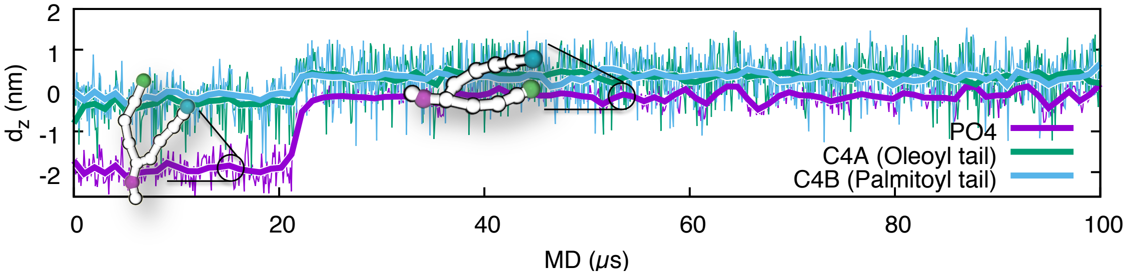


**Figure S8. Dynamics of the head group and two tails of a lipid as the head group moves up in the groove.** The result is from a 100 $\mu s$ CG MD trajectory of the CG model with a POPS/POPE/POPC mixed outer leaflet model which shows the locations of the phosphate group (PO4 bead) and two acyl chains (C4A and C4B beads) along the Z axis. The figure shows how a POPS lipid moves from the outer leaflet to the deep site of the groove, and keeps extending its hydrophobic acyl chains in the center of the membrane bilayer, therefore resulting in an approximate 90-degree rotation of the whole lipid.

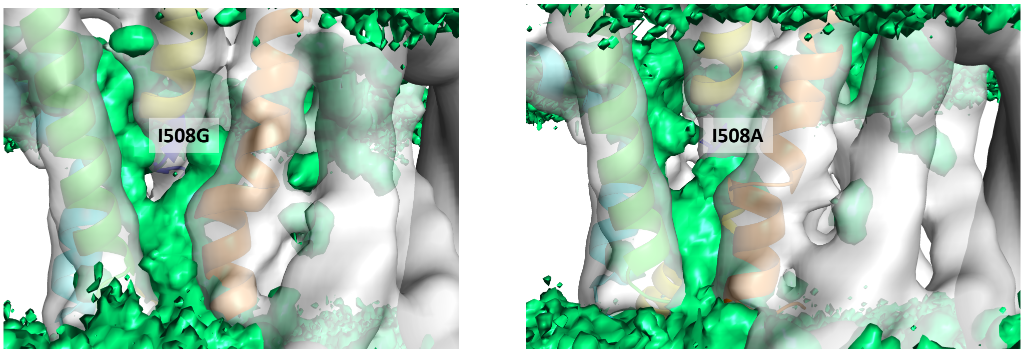


**Figure S9. Water-filled cavities in I508A and I508G mutants observed in all-atom simulations (AA5).** Average density map of the oxygen atoms in the water molecules (green surface). Each map was obtained from a 150 ns-long all-atom MD simulation. In both simulations, we observed rapid water exchange between the two water cavities in tens of nanosecond, suggesting that replacing Ile508 with alanine or glycine disrupts the seal between these two cavities.

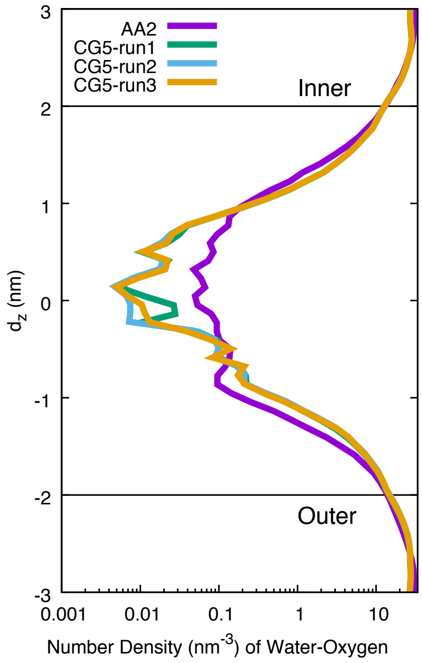


**Figure S10. Comparison of the water density profiles obtained from atomistic force field and coarse-grained models.** The densities from the Martini simulations have been corrected taking into account that a single coarse-grained water molecule represents four actual water molecules.

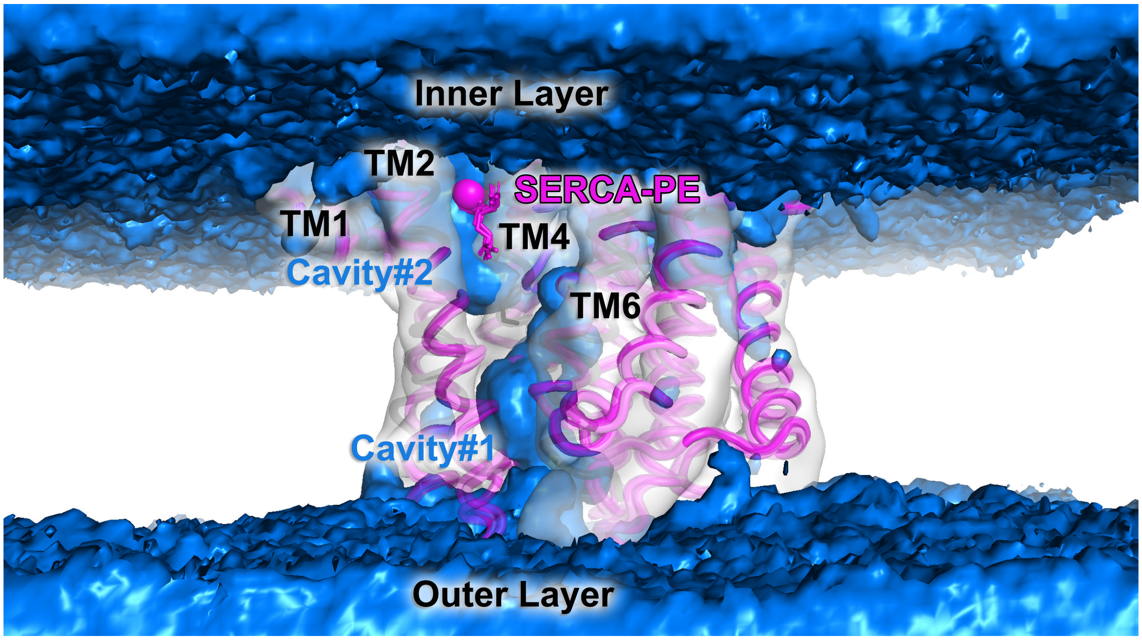


**Figure S11. Crystal structures of SERCA belong to P2 subfamily of P-type ATPases captured a PE molecule bound specifically in the same location that overlaps with the water-filled cavity on the inner cytosolic layer ('Cavity#2').** The average density map of the oxygen atoms of water molecule (blue surface) obtained from a 500 ns atomistic MD simulation with the pure POPS outer layer model (AA1 model). The cavity on the outer layer ('Cavity#1') overlaps with the TM2-TM4-TM6 groove and the cavity on the inner layer ('Cavity#2') is formed between TM1, TM2 and TM4. The structures of SERCA (PDB codes: 2AGV, 3AR4, 3W5C and 3W5D) are represented by magenta tubes. The phosphorus head groups of the bound PE molecules are shown in magenta spheres.

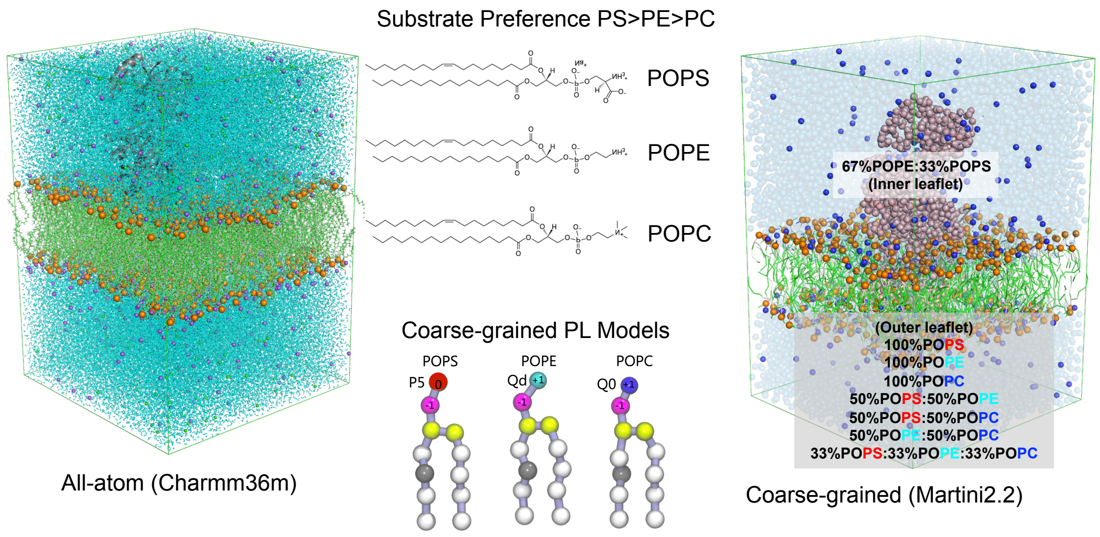


**Figure S12. Atomistic and coarse-grained modeling of the PI4P-activated E2P state of Drs2p.** The left and right panels show, respectively, the all-atom and coarse-grained systems that we simulated. The central panel shows the chemical structure of the three lipids, and their coarse-grained representations in the Martini model. We inserted the CryoEM structure of the PI4P-activated E2P state of the Drs2p-Cdc50p complex into an asymmetric lipid bilayer with 2POPE:1POPS in the inner leaflet and a varying ratio of POPS:POPE:POPC in the outer leaflet, resulting in seven different CG models and five atomistic models (Table 1).

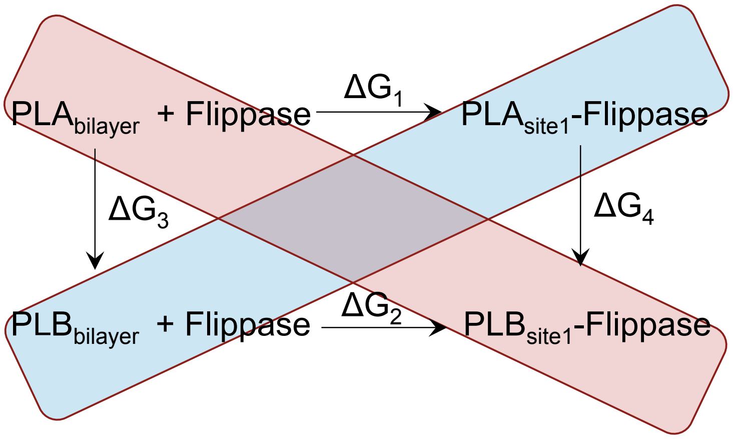


$$\Delta\Delta G_{PLA\rightarrow PLB} = \Delta G_2 - \Delta G_1 = \Delta G_4 - \Delta G_3$$

**Figure S13. Thermodynamic cycle used in the non-equilibrium alchemical free energy calculations.** We calculated $\Delta\Delta G_{PLA\rightarrow PLB}$ via the vertical branches ($\Delta G_3$ and $\Delta G_4$) using alchemical free energy calculations. Two vertical branches of the thermodynamic cycle were combined into a single transition by performing $PLA_{bilayer} \rightarrow PLB_{bilayer}$ and $PLB_{site} \rightarrow PLA_{site}$ transitions at the same time using the so-called 'double-system/single-box' setup.

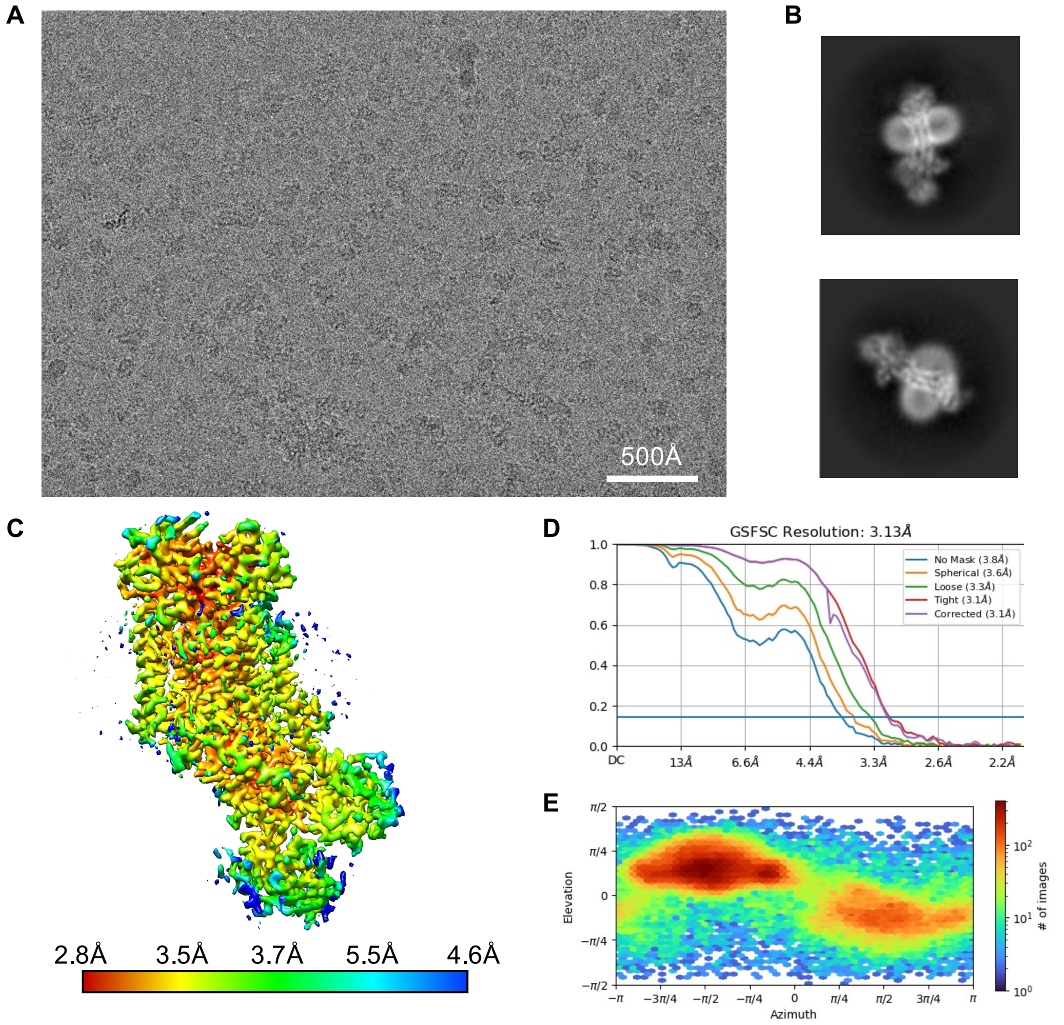


**Figure S14. Cryo-EM data and processing.** (A) Example micrograph collected at a defocus of -1.1 $\mu$m. (B) Example of good 2D class averages. (C) Final cryo-EM density colored by local resolution (calculated in cryoSPARC). (D) Fourier Shell Corellation (FSC) for the final volume (from cryoSPARC). (E) Angular distribution of the particles from the final refinement (from cryoSPARC).

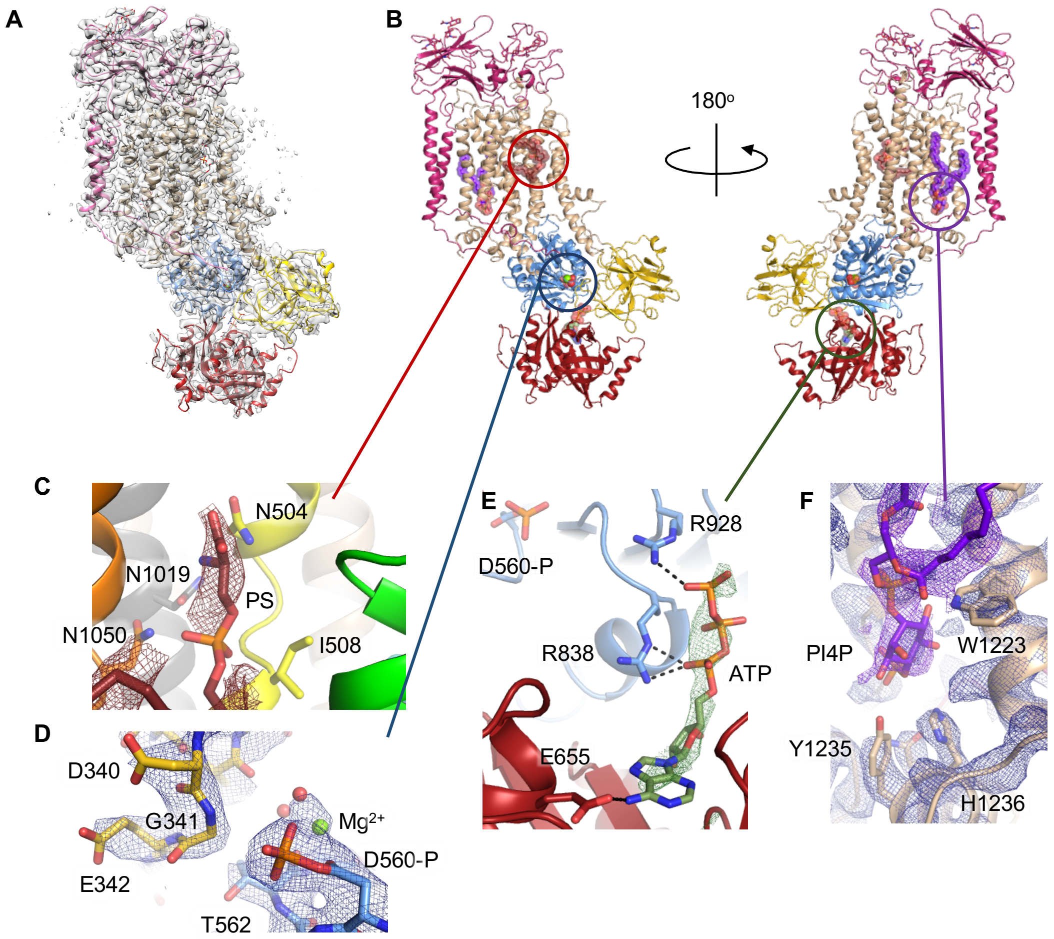


**Figure S15. Close-up of PS-bound and phoshorylated E2P-ATP state of Drs2p-Cdc50p.** (A) The cryo-EM density (at a contour level of 0.19) with the model fitted. For Drs2p the TM-domain is in tan, while the cytosolic domains, A-, N-, and P- are yellow, red, and blue, respectively. Cdc50p is shown in pink. (B) The model of the [PS]E2P-ATP state with ligands shown. Transport substrate PS is maroon (C), the phosphorylation site is indicated in blue (D), ATP is green (E), while the regulatory lipid PI4P is purple (F). For each ligand the density is shown as a coloured mesh, where the purple density of PI4P is at a lower contour level than the protein in panel F. The lower resolution of the PS density in panel C is likely caused by the transient nature of the PS-bound E2P-ATP state.

| | **[PS]E2P-ATP** (EMDB-13353) (PDB 7PEM) |
|---|---|
| **Data collection and processing** | |
| Magnification | 130,000x |
| Voltage (kV) | 300 |
| Microscope | Titan Krios G3 |
| Camera | Gatan K3 |
| Physical pixel size (Å/pix) | 0.66 |
| Electron exposure (e–/Å$^2$) | 60 |
| Defocus range (μm) | 0.3-2.3 |
| Number of movies | 8846 |
| Initial particle images (no.) | 3,029,402 |
| Final particle images (no.) | 91,415 |
| Symmetry imposed | C1 |
| Map resolution (Å) | 3.1 |
| FSC threshold | 0.143 |
| Map resolution range (Å) | 2.8-4.6 |
| **Refinement** | |
| Initial model used (PDB code) | 6ROJ |
| Model resolution (Å) | 3.3 |
| FSC threshold | 0.5 |
| Map sharpening B factor (Å$^2$) | -69.1 |
| Model composition | |
| Non-hydrogen atoms | 11620 |
| Protein residues | 1416 |
| Water | 2 |
| Ligands | 1 $Mg^{2+}$, 1POPS, 1 PI4P, 1 ATP, 6 NAG, 2 BMA |
| B factors (Å$^2$, min/max/mean) | |
| Protein | 20/126/61 |
| Ligand | 44/108/66 |
| Water | 45/49/47 |
| R.m.s. deviations | |
| Bond lengths (Å) | 0.002 |
| Bond angles (°) | 0.465 |
| Validation | |
| MolProbity score | 2.34 |
| Clashscore | 6.09 |
| Poor rotamers (%) | 6.14 |
| Ramachandran plot | |
| Favored (%) | 94.0 |
| Allowed (%) | 6.0 |
| Disallowed (%) | 0.0 |

**Figure S16. Cryo-EM data collection, refinement and validation statistics.**
